# [^123^I]Italia: A PARP-Directed Auger Electron-Emitting Agent for Targeted Radionuclide Therapy of Cancer

**DOI:** 10.64898/2026.03.13.711622

**Authors:** Sreelakshmi Unnikrishnan, Chiara Rua, Ge Li, Nerea Delgado Mayenco, Luis Hernandez Cano, Gülbahar Bozan, Ilias Patmanidis, Sangwani Simwaka, Ahmad Kurniawan, Wiktor Szymanski, Erik F.J. de Vries, Philip H. Elsinga, Ines Farinha Antunes, Gianluca Destro, Bart T. Cornelissen

**Author notes:** **1^st^ Author in training** contact details Sreelakshmi Unnikrishnan, MSc University Medical Center Groningen, University of Groningen Department of Nuclear Medicine and Molecular Imaging Hanzeplein 1, Groningen, The Netherlands. **To whom correspondence should be addressed:** Prof Bart Cornelissen, PhD, University Medical Center Groningen, University of Groningen Department of Nuclear Medicine and Molecular Imaging Hanzeplein 1, Groningen, The Netherlands, Dr. Gianluca Destro.

## Abstract

Poly(ADP-ribose) polymerase 1 (PARP1) is a central mediator of DNA damage repair and an established therapeutic target in homologous recombination-deficient cancers. Radiolabelled PARP inhibitors provide a strategy to deliver cytotoxic radiation directly to tumour DNA by exploiting PARP overexpression and trapping at sites of DNA damage. Here, we describe the design, radiosynthesis, and *in vitro* evaluation of [^123^I]Italia, a talazoparib-derived Auger electron-emitting agent for PARP-targeted radionuclide therapy. Stereochemically pure [^123^I]Italia, (8S,9R)-5-fluoro-8-(4-(iodo-^123^I)phenyl)-9-(1-methyl-1H-1,2,4-triazol-5-yl)-2,7,8,9-tetrahydro-3H-pyrido[4,3,2-de]phthalazin-3-one was synthesised in one step via copper-mediated iodo-deboronation, achieving activity yields >80% and molar activities >6.2 ± 3.1 GBq/µmol (n=8). UPLC analysis confirmed radiochemical purity >97%. Italia exhibited potent PARP1 inhibition (IC_50_ 0.48 nM) and in silico predicted binding affinity comparable to talazoparib. In a panel of PARP-expressing cancer cell lines, [^123^I]Italia demonstrated highest uptake at 60 min, PARP-selective uptake, predominant nuclear localisation (up to 60% of added activity) and chromatin association consistent with PARP trapping (up to 15% of total activity recorded). Uptake was reduced more than 50-fold by addition of an excess of any PARP inhibitor (e.g. olaparib, talazoparib, and rucaparib) and in PARP1 knockout cells, confirming target specificity.

Clonogenic assays showed a marked, added activity-dependent reduction in survival of PARP-expressing cells following a brief one-hour exposure, whereas PARP1-deficient cells were resistant. Collectively, these findings identify [^123^I]Italia as a promising PARP-targeted Auger electron-emitting theranostic candidate that warrants further *in vivo* evaluation.

## Introduction

Cells are persistently exposed to endogenous and exogenous factors that damage their DNA. Indeed, each cell in the human body experiences tens of thousands of DNA damage events daily from sources such as metabolic byproducts, oncogenic signalling, ionizing radiation, and chemical exposure.^1–4^ Although most damage is repaired, unrepaired damage accumulates, driving genomic instability and contributing to cancer development. Poly(ADP-ribose) polymerase (PARP) is a highly abundant and widely expressed enzyme that plays a central role in DNA damage repair and gene expression regulation. PARP serves as a first responder to DNA single-strand breaks, catalysing PARylation of itself (auto-PARylation) and other nuclear proteins, using NAD^+^ as the donor molecule.^5–8^ Auto-PARylation is necessary for PARP to be removed from the site of DNA damage, to allow further DNA repair machinery to eventually repair the break. The wider PARP family comprises 17 isoforms. Among these, PARP1 is recognised for its pivotal role in DNA damage repair, alongside its involvement in chromatin remodelling, transcription, and cell death signaling.^9–11^ PARP inhibitors competitively occupy the NAD^+^ binding site, blocking auto-PARylation and thus prevent the release of PARP from damaged DNA, effectively trapping the enzyme at the break site.^12–14^ PARP inhibitors are primarily used in homologous repair defective cancers, where blocked DNA repair leads to cell death, while normal, homologous recombinant-proficient cells survive, exemplifying synthetic lethality, which allows PARP inhibitors to target cancer cells with minimal harm to healthy tissue.^15–18^ Four PARP inhibitors, namely olaparib, rucaparib, niraparib, and talazoparib, have so far been approved by the FDA and EMA for clinical use in patients with homologous repair defective cancers.^19,20^

PARP inhibitors labelled with therapeutic radionuclides, on the other hand, selectively deliver ionizing radiation to tumour DNA, exploiting (1) the markedly higher PARP1 expression in cancer cells compared to normal tissues, (2) the trapping of PARP by PARP inhibitors to DNA, and (3) the relatively larger amount of PARP associated with DNA in tumour compared to normal tissue.^21,22^ Localisation of PARP inhibitors at PARP1 binding sites places the radionuclide in close proximity to DNA, making therapeutic efficacy dependent on the range of emitted radiation. Radionuclides that emit ultra-short-range radiation are therefore particularly advantageous. In this context, Auger electron-emitting radionuclides are highly attractive, as they deposit energy over nanometre-scale distances, inducing localised DNA damage. Upon PARP1 binding, the radiolabelled PARP inhibitor releases radiation that ultimately leads to tumour cell death.^23–25^

Over the past decade, multiple radiolabelled PARP inhibitors have been developed for imaging (PET and SPECT alike) and targeted radionuclide therapy, including radiolabelled versions of olaparib, rucaparib, talazoparib, and veliparib, employing a variety of radionuclides. PARP targeted radionuclide therapy has been explored using α-, β-, and Auger electron emitters, including ^77^Br, ^123^I, ^125^I, ^131^I, and ^211^At, incorporated into compounds structurally related to olaparib or rucaparib.^21,22,23,26,27^ However, despite these advances, several challenges remain, including limited selectivity among PARP family members, non-specific uptake, and suboptimal pharmacokinetic profiles.^28^

Given talazoparib’s high binding affinity for PARP1 and superior PARP-trapping capability, a talazoparib-based compound is expected to be particularly well suited for PARP-targeted radionuclide therapy, as these properties are critical for enhancing localised radiation DNA damage in tumour cells. On this basis, we hypothesised that a talazoparib-derived radio-analogue could represent a promising candidate for Auger electron-based PARP-targeted radionuclide therapy. While Bowden et al. reported the development of radiofluorinated [^18^F]talazoparib^29^, we now present a radioiodinated analogue, [^123^I]Italia, for Auger electron PARP-targeted radionuclide therapy (**Figure 1**). In this manuscript, we describe its synthesis, radiochemical properties, and in-vitro evaluation, supporting its potential as a clinically relevant targeted radionuclide therapy agent.

**Figure 1:**
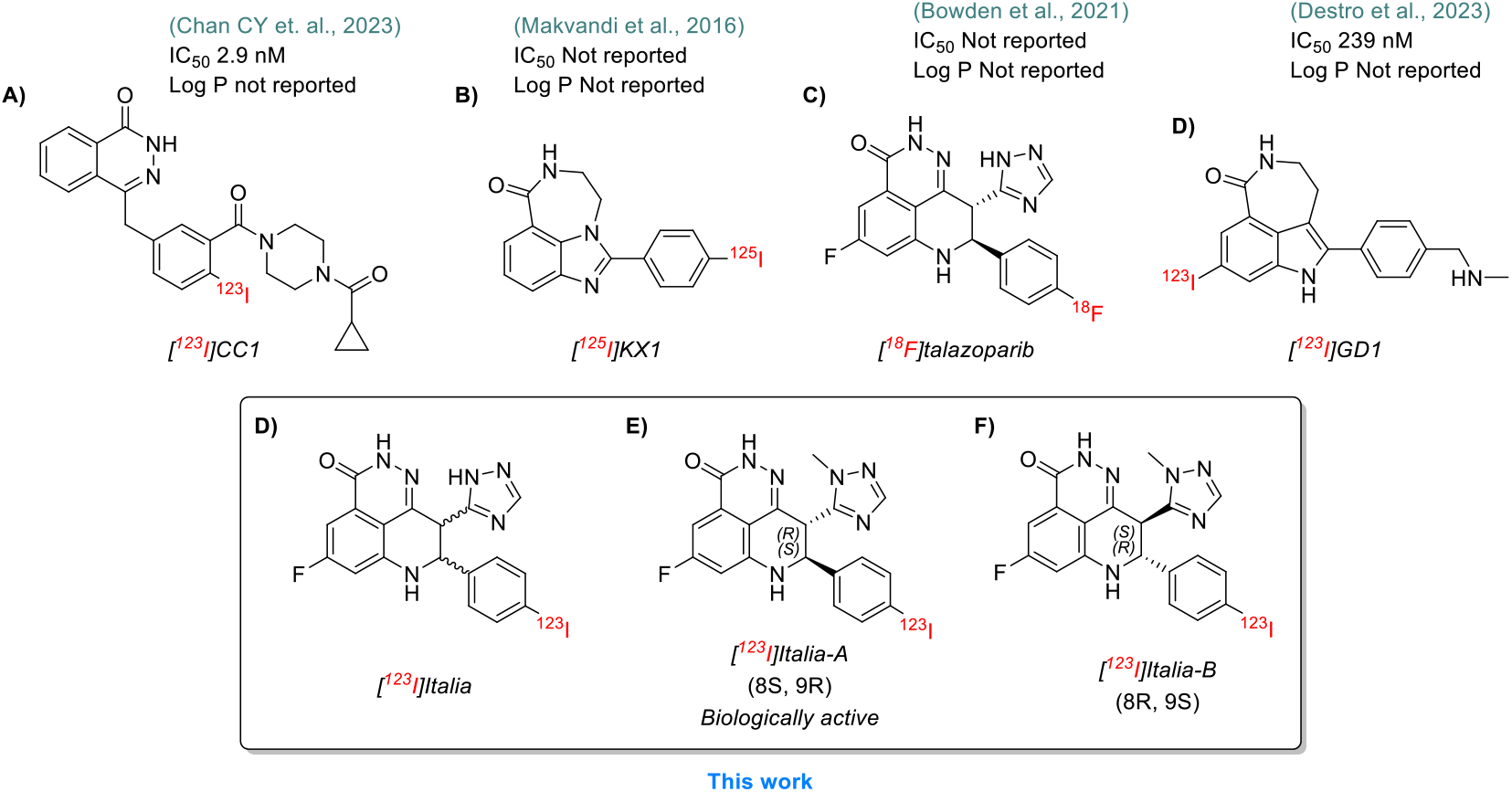
Examples of reported radiolabelled PARP imaging and radionuclide therapy agents.

**Figure 2:**
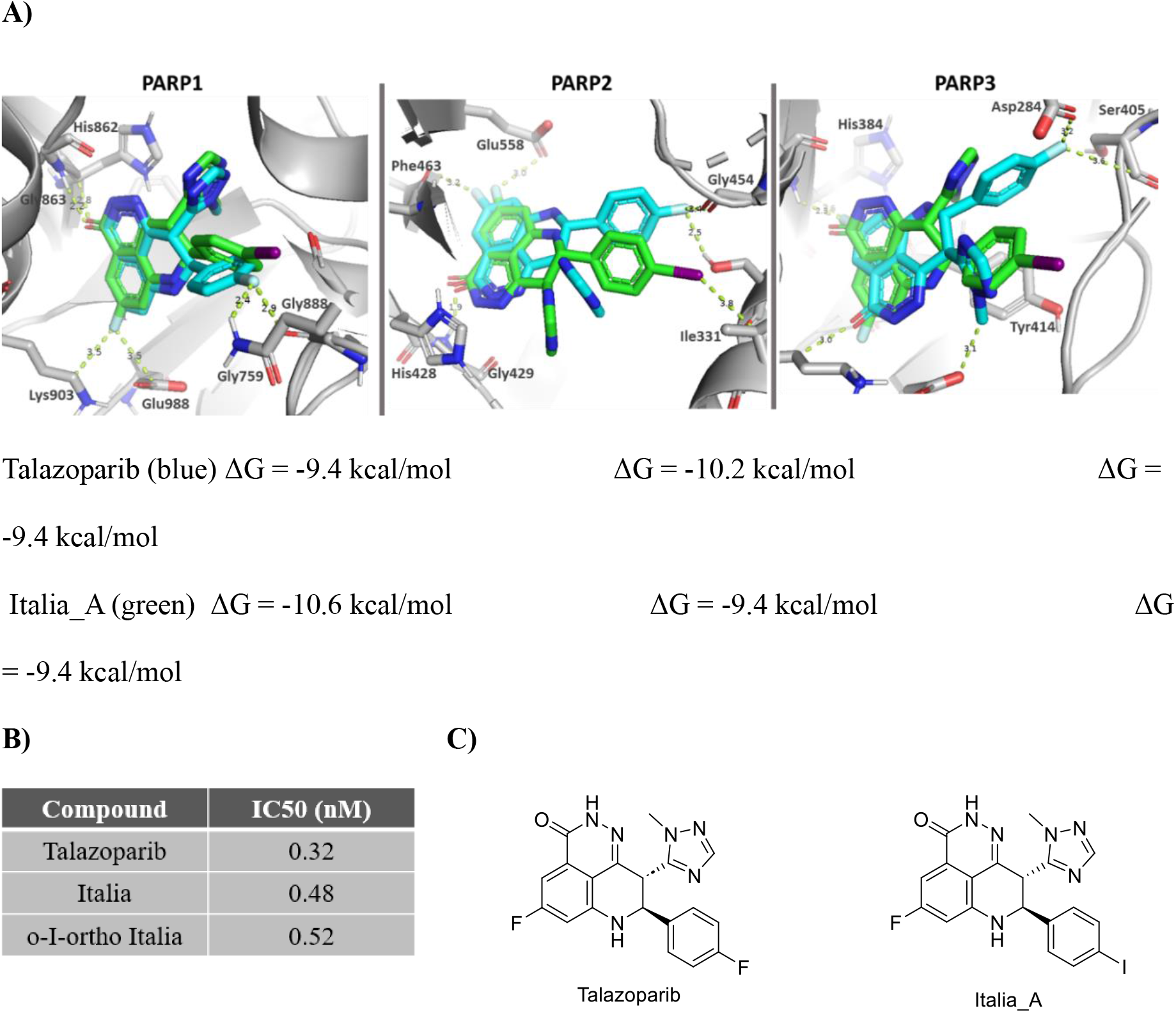
Molecular docking, enzymatic inhibition, and chemical structures of Italia compounds and talazoparib. **A)** Snapshots from the molecular docking study of talazoparib (blue) and Italia _A (green) to PARP1, PARP2, and PARP3, shows excellent overlap. **B)** Cell-free enzymatic inhibition of PARP1 by Italia (**8**), ortho-Italia (**10**) or talazoparib. IC_50_ = inhibitory concentration of 50% (in SI). **C)**Chemical structures of Italia_A and talazoparib.

## MATERIALS AND METHODS

### Precursor synthesis

General synthesis of iodinated talazoparib derivatives **8** (non-radiolabelled Italia), **9, 10**, boronic acid precursor **11**, and the synthesis of Italia was adapted from Bowden et al.^29^ [Cu(OAc)(phen)_2_]OAc was produced from copper(II)acetate and 1,10-phenanthroline under basic conditions, as previously described.^30^ Italia was obtained over 4 steps from commercially available compounds (Supplemental Scheme 1). Chiral separation of enantiomers of precursor **11** was performed using ChiralPak IB N5 column (30% MeCN/70% water/0.1% TFA).

### Molecular Docking Studies

The crystal structure of PARP1 in complex with talazoparib (PDB: 7KK3) was used as the reference for docking studies. PARP1, PARP2, and PARP3 structures (AlphaFold3) were aligned to the reference, and docking was performed using AutoDock Tools with default parameters and a grid centred on the crystallographic binding site. Italia (8*S*,9*R* enantiomer, Italia_A) was compared to talazoparib based on predicted binding energies and docking poses. Full details are available in the supplemental materials.

### Synthesis of [^123^I]Italia_A

**Scheme1:**
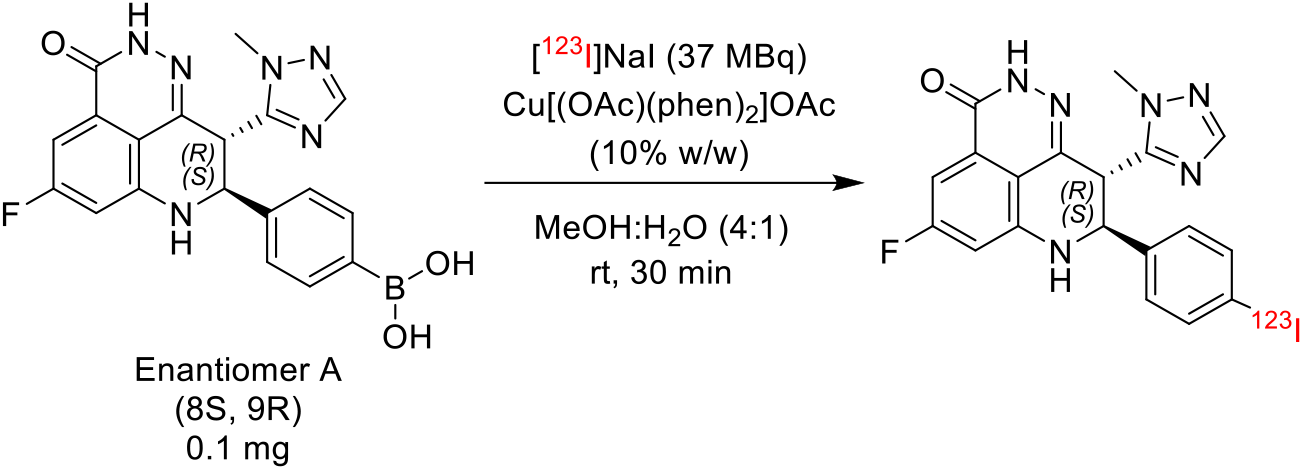
Radiosynthesis of [^123^I]Italia_A

Na[^123^I]I was obtained from GE Healthcare as a non-carrier-added formulation in 0.05 M NaOH (typically delivered as a 18.7±1.4 GBq/mL solution). Radiolabelling of [^123^I]Italia was performed in a 5 mL glass V-vial containing 0.5 mg boronic acid precursor (4-(5-fluoro-9-(1-methyl-1H-1,2,4-triazol-5-yl)-3-oxo-2,7,8,9-tetrahydro-3H-pyrido[4,3,2-de]phthalazin-8-yl)phenyl)boronic acid, **Scheme 1**). To this vial, 20 µL of [^123^I]NaI, diluted with 50 µL MeOH, and 0.05 mL of copper catalyst solution (Cu[(OAc)(phen)_2_]OAc 10% w/w in MeOH:H_2_O, 4:1), was added and the reaction was stirred at room temperature for 30 min. After completion, the reaction mixture was diluted to 1 mL with 50:50 MeCN/H_2_O and passed through a 0.2 µm Millex-25 hydrophobic PTFE vented filter. The filtrate (1 mL) was purified by semi-preparative reversed-phase HPLC, using a Luna Omega Polar-C18, (100 Å, 250 × 10 mm) column and 30% MeCN/H_2_O as the mobile phase at 2 mL/min, an Elite LaChrom VWR Hitachi L-2130 pump. Detection was achieved with a UV detector (Elite LaChrom VWR Hitachi L-2400) set at 254 nm and a Bicron Frisk-Tech radiation detector. The radioactive product fraction eluting at 24 minutes was collected and subsequently diluted with 20 mL of water and loaded onto an Oasis HLB cartridge (Waters, 30 mg), followed by rinsing the collecting vial with an additional 10 mL of water, which was also passed through the cartridge. The product was eluted with 0.5-1 mL of ethanol, evaporated at 80°C, and reconstituted in 0.5-1 mL of 10% DMSO/H_2_O to obtain the final formulation. Quality control was performed by UPLC using an ACQUITY UPLC BEH C18 column (1.7 µm, 2.1 × 50 mm) with 17% MeCN/H_2_O (containing 0.1% TFA) as the mobile phase at a flow rate of 1 mL/min with a retention time of 4.2 min. The radioiodination achieved radiochemical yields commonly ≥50%, with molar activities > 6.24 ± 3.09 GBq/µmol (n=8). LogP & LogD and human plasma stability of [^123^I]Italia were determined using the procedure detailed in the supplementary information. [^123^I]Italia_A was synthesised as above, but using the stereochemically purified boronic acid precursor (scheme 1).

### Radio-TLC

Radio-TLC analysis of [^123^I]Italia was performed on silica TLC plates. The TLC plate was co-spotted with a non-radioactive (“cold”) reference standard to unambiguously identify the expected R_f_ position. Plates were developed using 5% MeOH/DCM (v/v) as the mobile phase. After development, plates were dried, briefly visualised under UV, and the cold standard spot was marked in pencil. A small aliquot of [^123^I]Italia was then applied, allowed to dry, and exposed to a phosphor imaging screen. Radioactivity distribution was quantified using a Typhoon phosphor imager, and radiochemical purity was calculated by integrating activity in the product region relative to total activity using OptiQuant software.

### Cell free IC50

A commercially available assay was used to measure the catalytic activities of PARP1 (catalogue #80551) in a cell-free *in vitro* system in the presence of varying concentrations of established PARP inhibitors and non-radiolabelled Italia **8**, as well as ortho and meta variants **9** and **10**.

### Cell-free Pull-Down Studies

Single-strand break (ssBreak) biotinylated DNA (btDNA) constructs were incubated with His-Tagged Recombinant Human PARP (Medchem, HY-P74652) and [^123^I]Italia (100 kBq) in the presence or absence of olaparib (10 µM), or talazoparib (10 µM) for 1 hour at 30°C under constant agitation buffer (50 mM Tris-HCl pH 8, 10 mM MgCl_2_, 10 mM DTT and 10% Glycerol). Streptavidin-based pull-down assays using Thermo Scientific™ Pierce™ Streptavidin Agarose (according to the manufacturer’s instructions) was used to isolate btDNA and His-PARP complexes were isolated by anti-His immunoprecipitation using Protein G Agarose Resin 4 Rapid Run. ^123^I content in the fractions was measured using a gamma counter. Full details are available in the supplemental materials.^31,32^

### Cell culture

Human pancreatic adenocarcinoma cells (PSN-1), and human prostate cancer cells (PC-3) were purchased from ATCC; the murine breast cancer cell line 4T1 was a kind gift from prof. Nicola Sibson, Department of Oncology, University of Oxford. All three cell lines were cultured in RPMI medium. Human malignant glioma cells (U-87MG), human pancreatic cancer cells (MIA-PaCa2), and breast cancer cells (MDA-MB-231) were purchased from ATCC and maintained in high-glucose Dulbecco’s Modified Eagle’s Medium. All media were supplemented with 10% foetal bovine serum (Gibco), 2 mM L-glutamine, 100 units/mL penicillin, and 0.1 mg/mL streptomycin (Gibco). All cell cultures were maintained at 37°C in a humidified atmosphere containing 5% CO_2_. Cells were passaged and harvested using trypsin-EDTA solution and were utilised for no more than 20 passages following resuscitation from liquid nitrogen storage. Cell identity was confirmed by the provider and by short tandem repeat profiling, and regular testing was performed to ensure the absence of mycoplasma. PSN1^PARP1−/−^ knockout cells were generated through CRISPR-Cas9 (see supplementary information for additional methodology and PARP1 knockdown validation).

### *In vitro* Uptake, Efflux, and Specificity of [^123^I]Italia

To investigate PARP-mediated specific uptake, cells were harvested using trypsin-EDTA, seeded into 24-well plates (10^5^ cells per well) containing growth medium (1 mL), and allowed to adhere overnight. Cells were then washed and exposed to vehicle control, unlabelled PARP inhibitors or unlabelled Italia (10 µM) in 500 µL of growth medium for 60 minutes at 37°C. This was followed by [^123^I]Italia (10 µL, 10 kBq/well, n=3), and the cells were further incubated for 60 min. Medium was aspirated, the cells washed twice with phosphate-buffered saline (PBS), and cells were lysed by adding 0.1 N NaOH for 10 min at room temperature. The amount of ^123^I in the cell lysates was measured using an automated gamma counter (PerkinElmer). Cell numbers were determined with non-lysed cells using an automated cell counter (Countess).

To investigated tracer uptake kinetics, cells were exposed to [^123^I]Italia (10 kBq) at 37°C for various time intervals (15-180 min). The amount of ^123^I in the cell lysates was measured as described above. To investigate efflux of the tracer from the cells, cells were incubated with [^123^I]Italia for 60 min at 37°C, washed with PBS, and then supplied with fresh growth medium. The amount of ^123^I retained by the cells was subsequently measured at different time points as previously described.

### Cell fractionation - Trapping Experiment

Aliquots of 10^7^ suspended cells (PSN1, PSN1^PARP1−/−^) were exposed to 100 kBq of [^123^I]Italia for 60 min in a total volume of 500 µL, after 30 min exposure to an excess of olaparib (10 mM, 37°C) or vehicle control. Samples were placed on ice, followed by washing cells with ice-cold PBS. Nuclear soluble and chromatin-bound fractions were isolated using the Subcellular Protein Fractionation Kit for Cultured Cells (Thermo Scientific, 10455924) according to the manufacturer’s protocol. The levels of ^123^I in the different cell fractions, and supernatant were measured using a gamma counter.

### PARP levels associated with chromatin - Western Blot

Suspended cells (5 × 10^6^, PSN1) were exposed to 200 kBq/mL [^123^I]Italia as above. Chromatin and nuclear soluble fractions were isolated, and proteins were extracted using the Subcellular Protein Fractionation Kit for Cultured Cells (Thermo Scientific, 10455924) supplemented with protease inhibitors, 40 µg of protein were analysed by SDS-PAGE and transferred to PVDF membranes. PARP1 was detected by immunoblotting using specific primary and HRP-conjugated secondary antibodies, followed by chemiluminescent detection and quantification using Fiji/ImageJ.

### Colony Formation Assay

Aliquots containing 10^5^ cells (PSN1, 4T1, U-87MG, PC3, MIA-PaCa2, PSN1^PARP1−/−^, or PSN1^CAS9^ cells) were suspended in 1 mL growth medium in Eppendorf tubes and exposed to increasing concentrations of [^123^I]Italia (0-500 kBq) for 60 min at 37°C. From each tube 10 µL was added into 6-well plates, and growth medium was added to a final volume of 2 mL per well. After one to two weeks depending on the proliferation rate of each cell line, cell colonies (defined as clusters of >50 cells) were fixed and stained using crystal violet solution (1 mg/L in a 1:1 water-methanol mixture) and enumerated.

### Statistical Analysis

All experiments were performed at least in triplicate. Statistical analysis was performed using GraphPad Prism version 9 or higher (GraphPad Software). Data were assessed for normality and analysed using one-way or two-way ANOVA, as appropriate. Significance was reached at a P value of less than 0.05. Results are presented as mean ± SD unless otherwise specified.

## RESULTS

### Italia: A Selective and Potent PARP Inhibitor

Italia_A (8*S*,9*R*), which shares a structural scaffold with talazoparib (**Figure 1**; Supplementary Figure S2), fits well within the NAD^+^ binding pocket of PARP1, PARP2, and PARP3, closely resembling the binding mode of talazoparib. Notably, Italia achieved a higher docking score than talazoparib. While docking scores do not always directly correlate with binding affinity and may be influenced by lipophilicity, the results suggest that iodine substitution does not disrupt the original fluorine-binding position within the PARP1 pocket, and highlighting its potential for further development. Italia demonstrated potent PARP inhibitory activity, with an in-house determined cell-free IC_50_ of 0.48 nM, despite using a racemic mixture of two enantiomers (**Figure 1B**; Supplementary Figure S1). This value was comparable to that of talazoparib measured under the same assay conditions (0.32 nM).

### Radiosynthesis of [^123^I]Italia

[^123^I]Italia (A (8*S*,9*R*), B (8*R*, 9*S*), and racemic mixture) was synthesised from the relevant boronic acid precursor **11** via copper-mediated radioiodination using Cu[(OAc)(phen)_2_]OAc as the catalyst (**Scheme 1**).^30^ [^123^I]Italia was produced in good radiochemical yield and with high molar activity (A_m_). Radiochemical conversions exceeding 80%, RCY of ~50% and a molar activity >6.24 ± 3.09 GBq/µmol (n=8) were obtained after a 2-hour synthesis. Product identity was confirmed by co-elution with the non-radioactive Italia reference standard (Table1; entry 5).

Reaction conditions were optimised by varying precursor loading (**Table 1**). Reaction times of 30 min resulted in higher radiochemical conversion (RCC) compared with 10- and 20-minute reaction times (entry 3). Radioiodination proceeded at room temperature. High reaction temperatures (60°C, 80°C) led to reduced radiochemical yield (RCY), likely due to product degradation (Table 1, entries 4-7). To minimise precursor consumption while maintaining acceptable radiochemical yield, the lowest effective precursor loading (0.1 mg) was selected (Table 1, entry 11). Preliminary optimisation studies were performed using the racemic precursor mixture, which was subsequently extended to the radioiodination of enantiomers A and B separately (Table 1, entries 9-12).

**Table 1:**
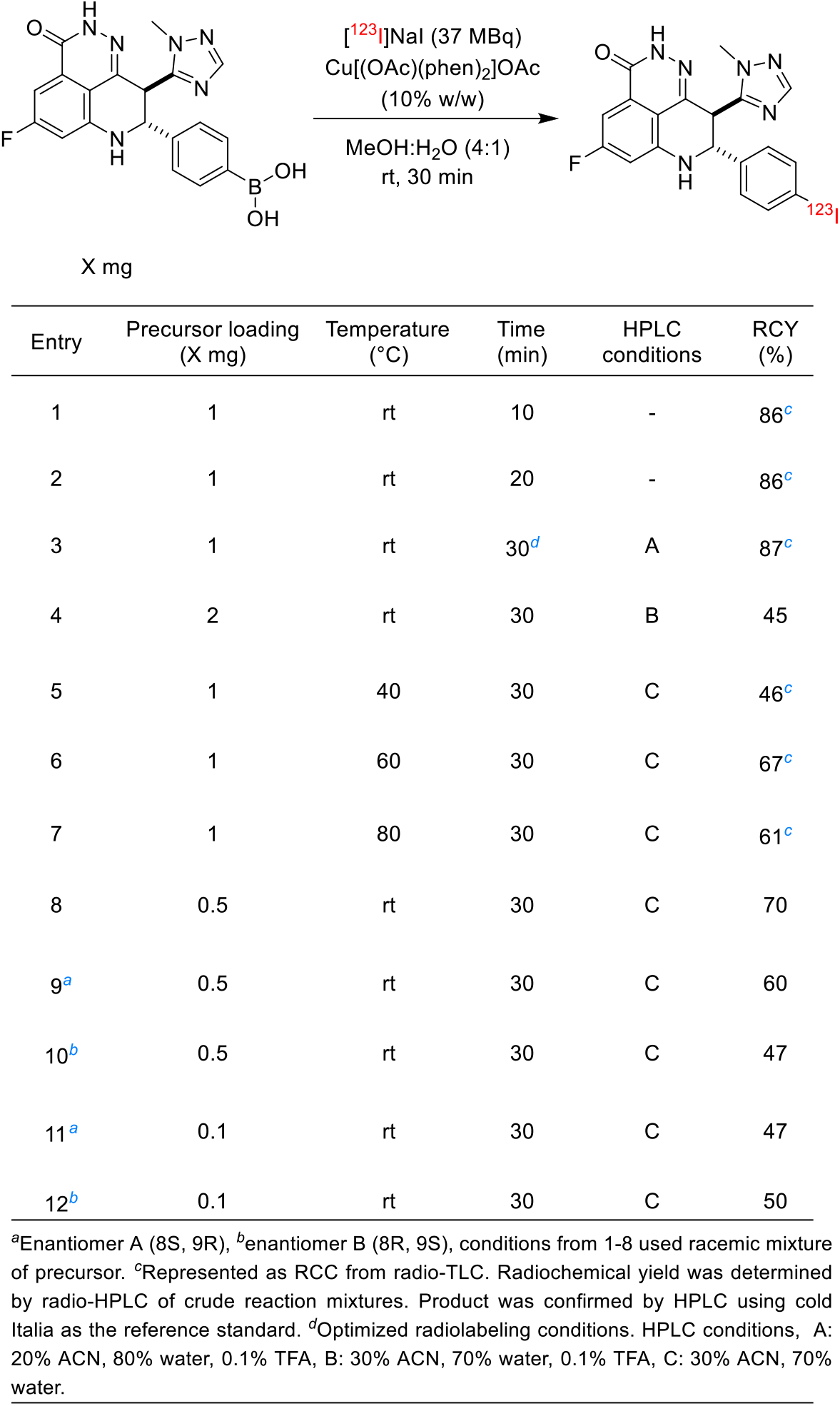
Optimisation studies of radiosynthesis of [^123^I]Italia by copper-assisted iododeboronation.

The lipophilicity assessment revealed a logP of 1.15 ± 0.28 and logD of 1.5 ± 0.34, indicating that [^123^I]Italia exhibits moderate lipophilicity. This balance suggests good membrane permeability while maintaining sufficient aqueous solubility for biological applications.

### *In Vitro* [^123^I]Italia Uptake in Cancer Cells is PARP-Selective

[^123^I]Italia was taken up in all PARP-expressing cells within minutes, peaking after 60 min (**Figure 4A, in SI, figure S8**). [^123^I]Italia exhibited a slow efflux, a decrease within the first 2 hours, and remained stable for up to 10 hours (**Figure 4B**).

[^123^I]Italia was taken up selectively in PSN-1, MDA-MB-231, U-87MG, and 4T1 cells, with an uptake up to 6 mBq/cell (**Figure 4C &** SI figure S7), but significantly less so in PSN-1^PARP1−/−^ knockout cells. One stereoisomer, [^123^I]Italia_A, exhibited far higher uptake than [^123^I]Italia_B, or the racemic [^123^I]Italia mixture (p<0.0001) (**Figure 4C)**. Addition of an excess of one of four structurally related or unrelated non-radiolabelled PARP inhibitors significantly reduced the cell-associated amount of [^123^I]Italia_A in all cell lines (p<0.0001), suggesting PARP-selective binding of [^123^I]Italia_A (**Figure 4D)**.

The racemic [^123^I]Italia displayed intermediate uptake, whereas [^123^I]Italia_B showed far lower uptake compared to the A isomer. All three formulations were competitively blocked by talazoparib. (**Figure 5A**). Fractionation of PSN-1 cells revealed predominant nuclear localisation of [^123^I]Italia_A, with approximately 50-60% of total recorded activity detected in the nuclear soluble fraction and 10-15% in the chromatin associated fraction. In contrast, PARP1 knock-out cells demonstrated significantly less nuclear accumulation and a relative increase in the membrane-associated fraction. Co-exposure to an excess of talazoparib significantly reduced tracer accumulation in the chromatin and nuclear fraction (p<0.0001) (**Figure 5C-F**). Exposure of cells to [^123^I]Italia_A increased PARP levels in the chromatin-bound fraction, consistent with PARP trapping on DNA. This effect was significantly reduced upon co-treatment with olaparib (10 µM), indicating competitive binding at PARP’s catalytic site. These findings confirm target-specific PARP engagement and chromatin association (**Figure 5B**).

### Pull Down

Pull down experiments were performed to evaluate the association of [^123^I]Italia_A with PARP and damaged DNA. PARP pull-down (**Figure 3A**) showed that [^123^I]Italia_A displays similar binding signal for PARP, whether associated with synthetically damaged DNA or not. Blocking of [^123^I]Italia_A binding to PARP by an excess of talazoparib indicated specific binding to PARP.

**Figure 3:**
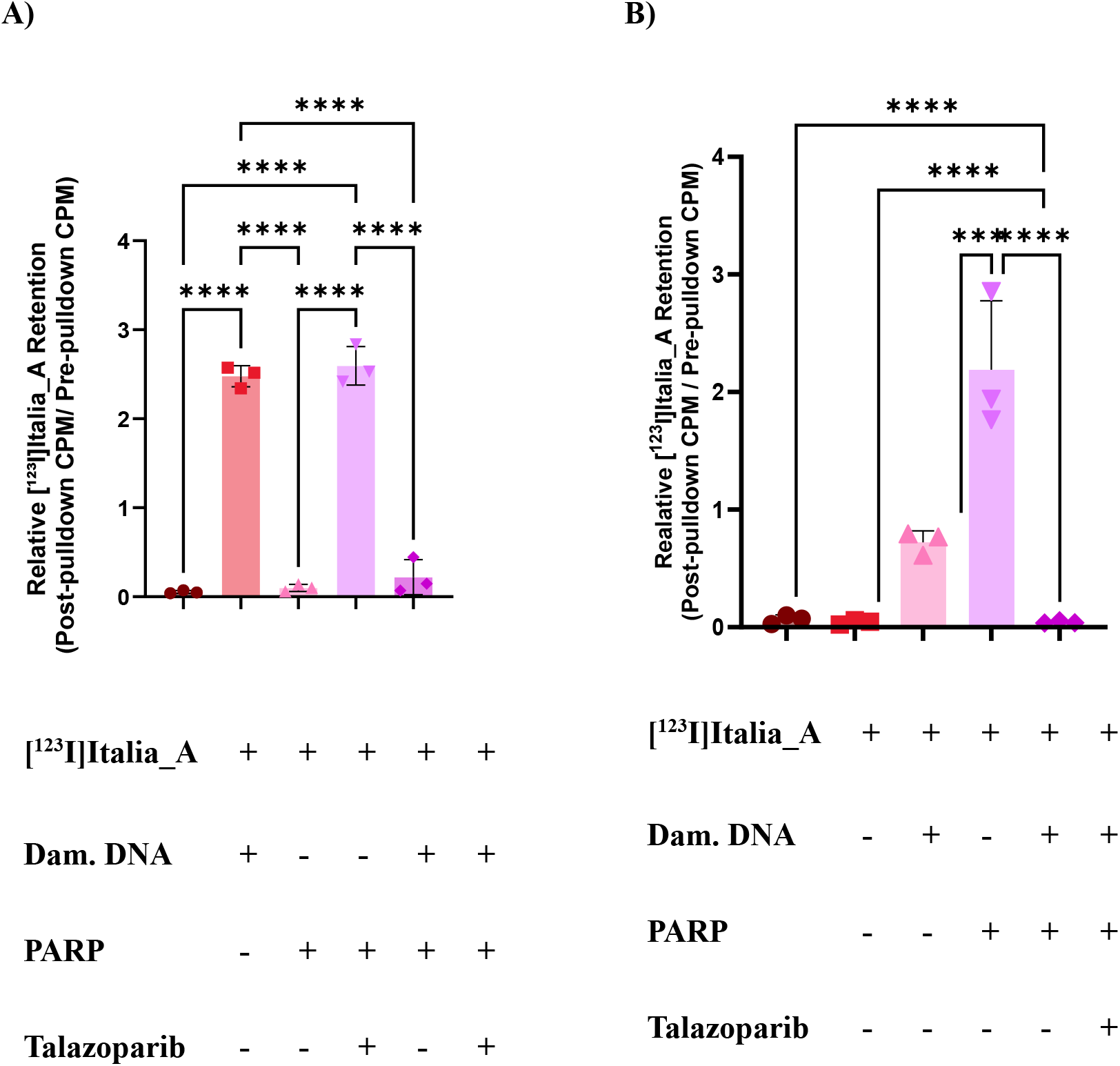
Pulldown assay demonstrating binding of [^123^I]Italia_A to PARP and damaged DNA in the presence or absence of talazoparib. **A**) Pull down from PARP. **B**) Pulldown from DNA.

**Figure 4:**
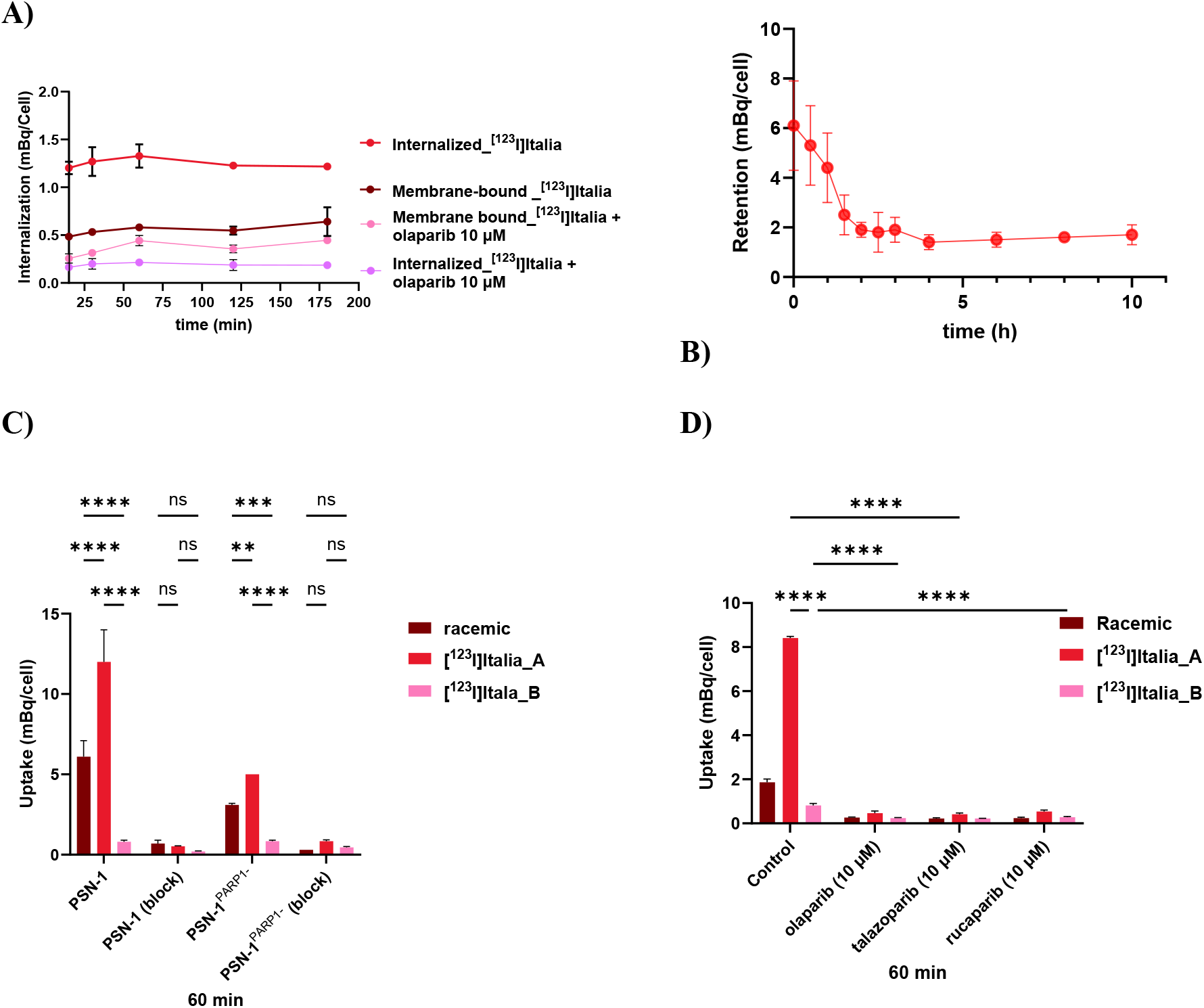
**A)** Uptake of [^123^I]Italia_A in PSN-1 cells at different time points. **B)** Efflux of [^123^I]Italia_A in PSN1 cells. **C)** Comparison of **u**ptake and blocking of [^123^I]Italia_A, B and racemic mix. in PSN-1, cas1, KO cells after 1 hour exposure. **D)** Comparison of **b**locking of [^123^I]Italia_A, B and racemic mix. uptake in PSN-1 cells by panel of PARP inhibitors.

**Figure 5:**
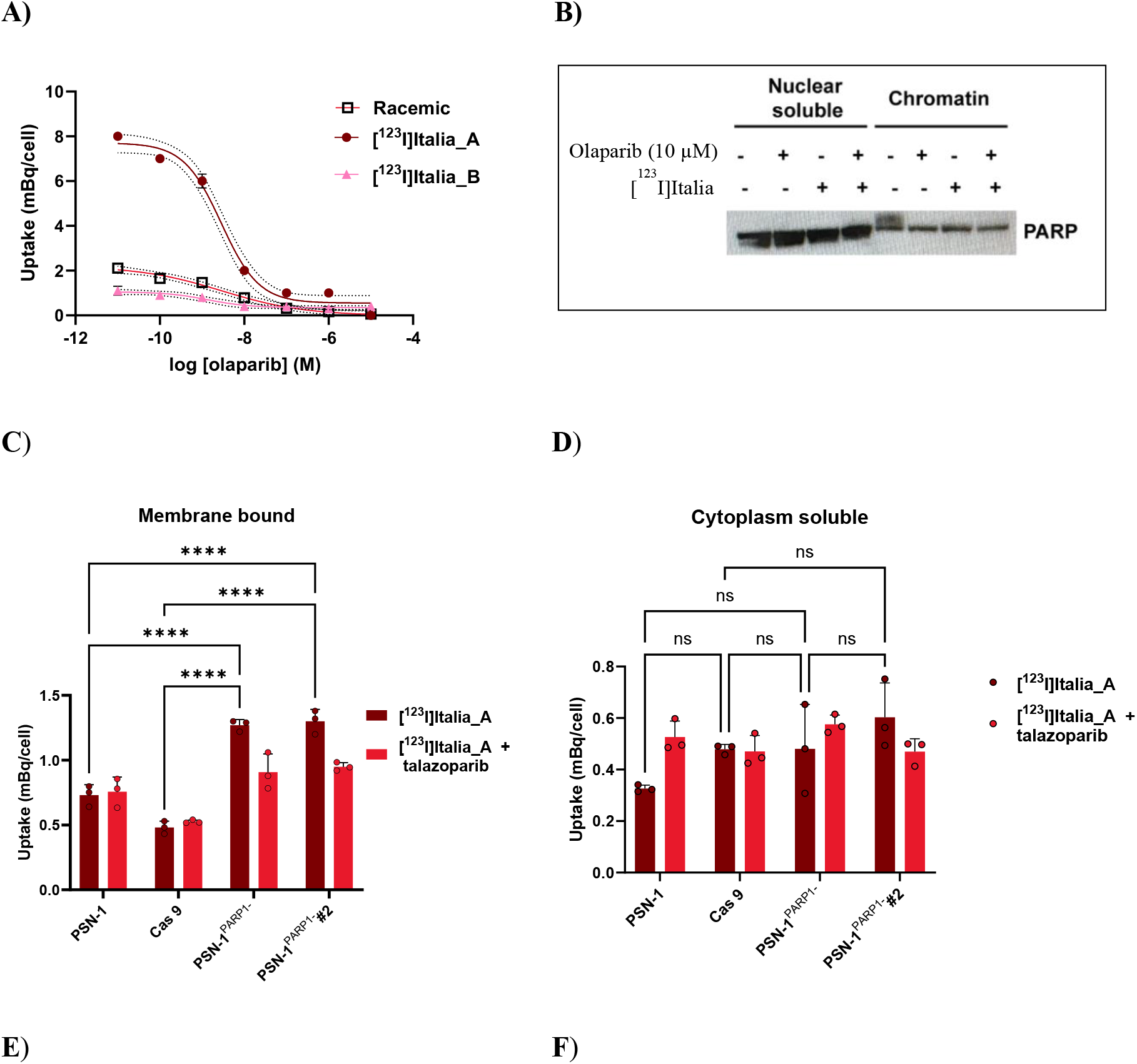
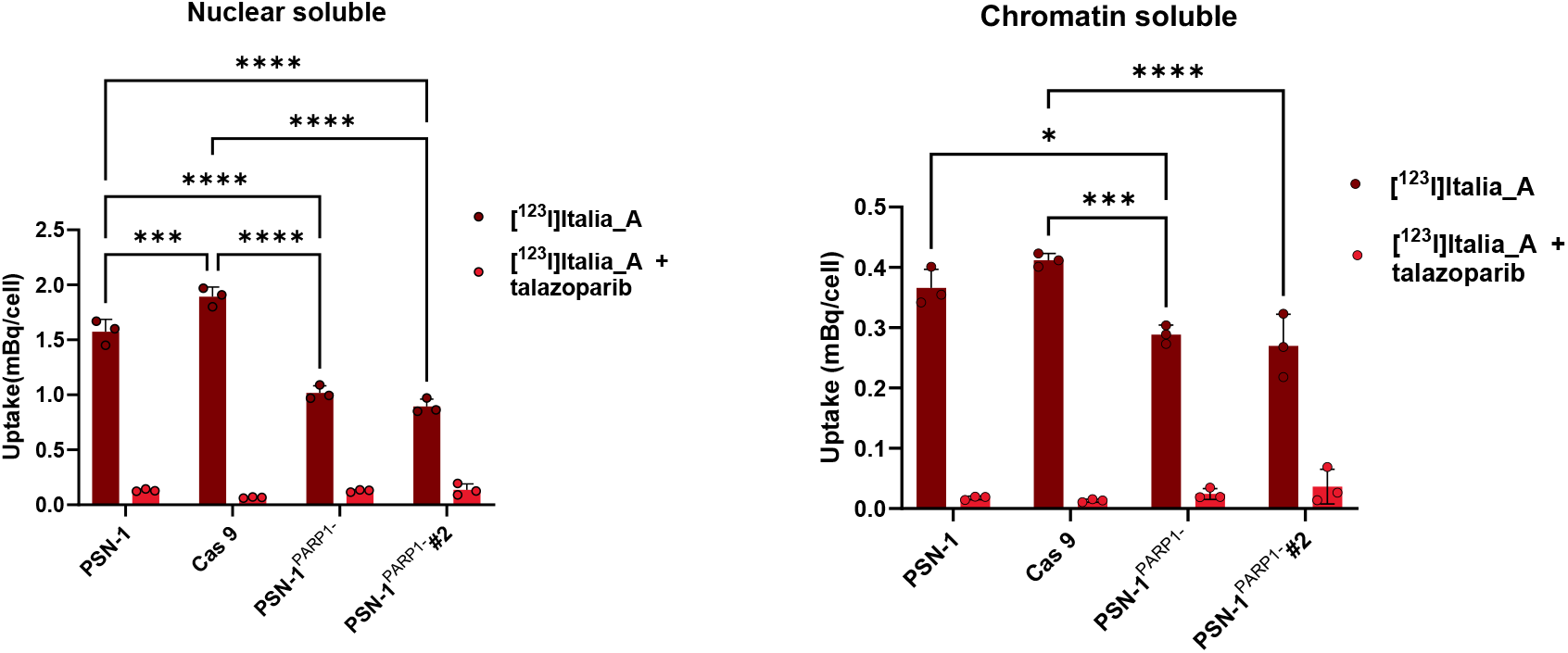
**A)** IC_50_ in PSN-1 cells (IC_50_: 2.85 nM, 95%CI: 1.98 nM-4.08 nM, R squared: 0.984 (of [^123^I]Italia), IC_50_: 1.36 nM, 95%CI: 0.5 nM −3.67 nM, R squared: 0.899 of [^123^I]Italia_B), IC50: 2.96 nM, 95%CI: 1.1 nM-7.9 nM, R^2^: 0.970 (of racemic mix.). **B**) Cell fractionation of [^123^I]Italia_A in PSN-1 cells: western blot. **C)** Cell fractionation of [^123^I]Italia_A in different cell lines. In membrane bound fraction. **D)** Cytoplasm soluble fraction. **E)** Chromatin soluble fraction. **F)** Nuclear soluble fraction.

Pull-down from biotinylated DNA demonstrated association of [^123^I]Italia_A with DNA in the presence of PARP (**Figure 3B**). Conversely, minimal binding was observed in the absence of PARP, of following co-incubation with an excess of talazoparib, confirming PARP-specific association with DNA.

### *In vitro* therapy: Clonogenic Survival

Clonogenic survival assays were performed to evaluate the radiotoxic effects of [^123^I]Italia enantiomers (**Figure 6A-B**). following a 1 h exposure of PSN-1, U-87MG, PSN-1^*PARP1*−/−^, PC-3, and MIA-PaCa2 cells. Brief exposure of PARP-expressing cancer cells to [^123^I]Italia_A significantly reduced their clonogenic survival, but not PARP1 knockout cells (**Figure 6A**). Clonogenic survival of cells was significantly reduced by exposure to [^123^I]Italia from added activities as small as 10 Bq. In PSN-1 cells, both A and B enantiomers decreased survival in a dose-dependent manner, but the A enantiomer showed significantly greater potency.

**Figure 6:**
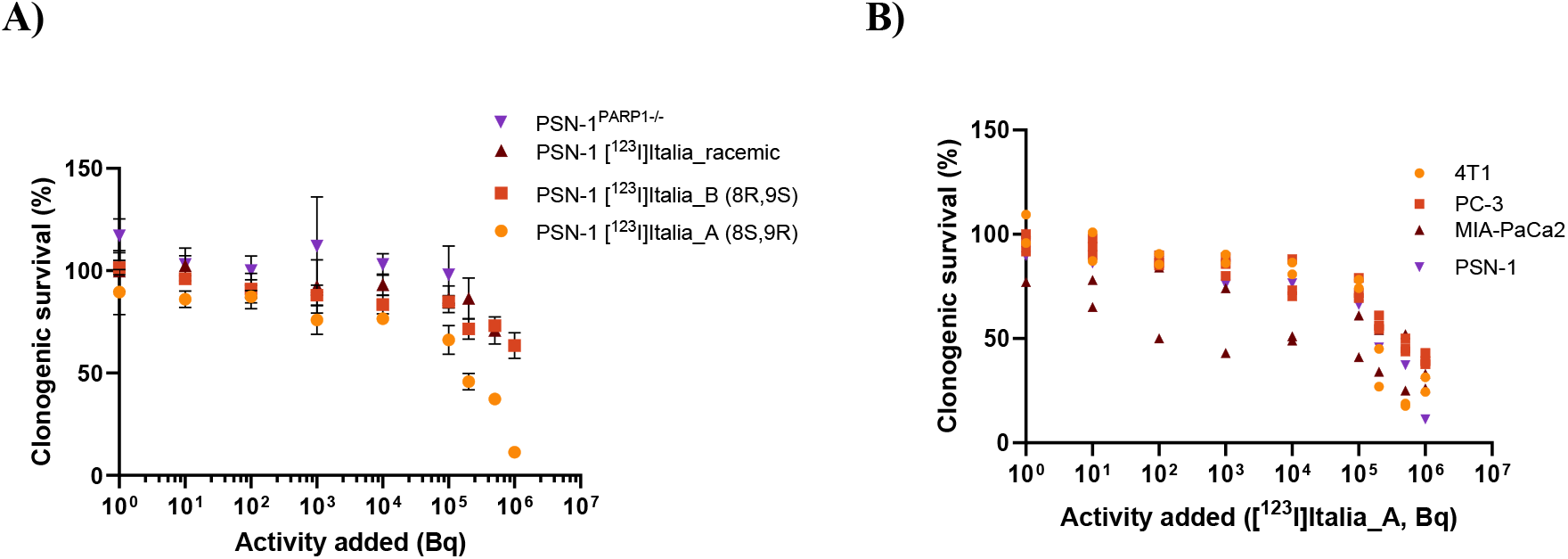
**A)** Clonogenic survival of PSN-1 and PARP1 knock out cells (PSN-1^PARP1−/−^) after exposure of cells for 1h to [^123^I]Italia_rac, and the separated enantiomers of [^123^I]Italia_A or _B (0-500 kBq in 2 mL of growth medium). **B)** Clonogenic survival of various cell lines after exposure of cells for 1h to [^123^I]Italia_A (0-500 kBq in 2mL of growth medium).

## Discussion

A variety of radiolabelled PARP inhibitors have been developed as potential PET and SPECT tumour imaging agents and targeted radionuclide therapy agents over the past few years. Most of these compounds are based on structural analogues of the clinically approved PARP inhibitors olaparib, rucaparib, and talazoparib, which have been modified to incorporate radioactive isotopes for molecular imaging applications.^33^ Talazoparib is regarded as the most potent PARP inhibitor and the most effective PARP-trapping agent currently in use.^34–36^ Its strong and sustained interaction with the PARP1-DNA complex suggests that radiolabelled derivatives may achieve prolonged retention at DNA damage sites, enhancing therapeutic efficacy. Talazoparib contains two chiral centres, giving rise to four possible stereoisomers. Among them, the anti-configured enantiomers were generated during synthesis, with the (8*S*,9*R*)-isomer (talazoparib) exhibiting high PARP inhibitory activity, whereas its enantiomer [(8*R*,9*S*), LT-674] and the remaining diastereomers are markedly less active.^37^

Molecular docking studies demonstrated that Italia_A, which retains the talazoparib scaffold, fits well in within NAD^+^ binding pocket of PARP1-3, and closely mimics talazoparib’s binding mode. Importantly, Italia achieved a more favourable docking score toward PARP1 than talazoparib and compares favourably with other clinically used PARP inhibitors such as olaparib, niraparib, and rucaparib.^38–40^

In this study, we developed an efficient one step radiolabelling process to synthesise stereochemically pure [^123^I]Italia from the corresponding boronic acid precursor. Conversely, previously reported [^18^F]talazoparib^29^ synthesis requires multiple steps, including deprotection and chiral separation, to afford stereochemically pure product. Employing a boronic acid precursor reduces the risk of hydrolytic degradation associated with boronic esters and eliminates concerns related to residual metal contamination and toxicity inherent to organotin precursors.^22,41,42,43^ The reaction proceeded efficiently with only a 40-fold less amount of precursor, in contrast to [^123^I]CC1, which required higher precursor amounts to achieve comparable radiochemical conversion. Apparent variations in molar activity were attributable to analytical limitations, as chromatographic sensitivity decreases below the limit of quantification (~1 mg/L) and detection (~0.4 mg/L), where baseline interference affects accuracy. Consequently, molar activity was conservatively calculated using and LOQ of 1 mg/L, with the true value likely much higher. Log P and log D values of [^123^I]Italia reflect its moderate lipophilicity are in good agreement with the reported log P value of talazoparib (1.6), enabling efficient membrane permeability while preserving sufficient aqueous solubility.^44^

We further focused our studies evaluating the *in-vitro* properties of [^123^I]Italia isomers on variety of PARP expressing cancer cell lines. Initial studies were conducted using racemic [^123^I]Italia and demonstrated approximately ten-fold higher PARP-specific uptake and internalisation compared with the previously reported [^123^I]CC1 in the same cell lines under the same conditions.^21^ While several PARP-targeted radiotracers, including [^123^I]MAPi and [^123^I]CC1, have demonstrated PARP-dependent uptake and cytotoxicity through blocking and knockout studies, direct biochemical evidence of catalytic-site engagement and PARP-dependent DNA association remains limited. We still observed residual uptake that could be blocked in PARP1 knockout cell lines. This suggests that [^123^I]Italia may not exclusively bind to PARP1. Furthermore, docking studies demonstrates that Italia_A binds not only PARP1, but also to PARP2 and PARP3 (Supplemental **figure S3**) which is in strong agreement with these observations. Additionally, genetic deletion of PARP1 could lead to compensatory upregulation of other PARP isoforms, which may contribute to the observed signal. Cell-free pull-down data presented here provide additional mechanistic validation that [^123^I]Italia_A binds selectively to PARP and is recruited to damaged chromatin in a trapping-dependent manner. These findings align with previously reported behaviour of high-affinity PARP inhibitors, particularly talazoparib, which is known to strongly stabilise PARP-DNA complexes.^21^

Consistent with previous reports describing distinct biologically active and less active enantiomers of PARP inhibitors, we investigated whether stereochemistry influences the biological performance of Italia. Marked stereochemical differences were observed, with [^123^I]Italia_A exhibiting significantly higher cellular uptake and functional activity than [^123^I]Italia_B, while the racemic mixture showed intermediate effects, consistent with the enantioselective behaviour described by Bowden et al. for [^18^F]talazoparib. Additionally, fractionation studies showed increased chromatin-bound PARP after treatment with [^123^I]Italia_A, an effect reduced by olaparib, confirming catalytic-site competition and PARP trapping. This retention is consistent with the strong trapping capacity of talazoparib-derived scaffolds and is particularly beneficial for Auger emitters such as iodine-123, which require close proximity to DNA for maximal cytotoxicity. [^123^I]Italia (A, B and racemic) binding was competitively inhibited by olaparib in a concentration-dependent manner, yielding IC_50_ values of 2.9 nM, 1.4 nM, 2.9 nM, respectively, and when inhibited with talazoparib yielded an IC50 of 0.53 nM against *rac*-[^123^I]Italia which is comparable to those reported for [^18^F]talazoparib by Bowden et al. (14 (talazoparib) and 24 nM (olaparib)).

The therapeutic rationale of ^123^I-labeled PARP inhibitors is based on PARP trapping, which keeps the radiolabelled compound bound at sites of DNA damage and places the Auger emitter in very close proximity to chromatin. Compared with previously reported data for [^123^I]CC1, [^123^I]Italia_A induces a pronounced and dose-dependent decrease in survival across multiple PARP-expressing cancer cell lines, yet at lower radioactivity doses.

This study is limited to *in vitro* evaluation, and further investigation into tumour-bearing animal models is necessary to assess metabolic stability, pharmacokinetics, metabolism, tumour uptake, normal tissue distribution, and therapeutic index *in vivo*. The dual imaging and therapeutic properties of ^123^I position [^123^I]Italia_A as a promising theranostic candidate. The combination of potent PARP engagement, and demonstrated Auger electron-mediated cytotoxicity suggests that [^123^I]Italia_A may serve as a true theranostic agent.

## Conclusion

Stereochemically pure [^123^I]Italia was successfully radiosynthesised. [^123^I]Italia integrates strong PARP targeting, efficient nuclear localisation, and Auger electron-mediated cytotoxicity with the imaging capability of iodine-123, supporting its role as a promising theranostic candidate. These promising *in vitro* results support further *in vivo* evaluation and preclinical development.

