## Supplemental data for "[^123^I]Italia: A PARP-Directed Auger Electron-Emitting Agent for Targeted Radionuclide Therapy of Cancer"

#### Contents

1. General .......... S2

2. Synthesis: .......... S3

3. Synthesis of compounds 8, 9, 10 and 11: .......... S3

3.1 5-fluoro-8-(4-iodophenyl)-9-(1-methyl-1H-1,2,4-triazol-5-yl)-2,7,8,9-tetrahydro-3H-pyrido[4,3,2-de]phthalazin-3-one (8) .......... S3

3.2 5-fluoro-8-(3-iodophenyl)-9-(1-methyl-1H-1,2,4-triazol-5-yl)-2,7,8,9-tetrahydro-3H-pyrido[4,3,2-de]phthalazin-3-one (9) .......... S4

3.3 5-fluoro-8-(2-iodophenyl)-9-(1-methyl-1H-1,2,4-triazol-5-yl)-2,7,8,9-tetrahydro-3H-pyrido[4,3,2-de]phthalazin-3-one (10) .......... S5

3.4 (4-(5-fluoro-9-(1-methyl-1H-1,2,4-triazol-5-yl)-3-oxo-2,7,8,9-tetrahydro-3H-pyrido[4,3,2-de]phthalazin-8-yl)phenyl)boronic acid (11) .......... S5

4. NMR Spectra .......... S6

4.1 $^{1}$HNMR Compound 8 .......... S6

4.2 $^{1}$HNMR Compound 9 .......... S7

4.3 1HNMR Compound 10 .......... S8

4.4 $^{1}$HNMR Compound 11 .......... S8

4.5 $^{13}$CNMR Compound 11 .......... S9

5. Chiral HPLC separation of Italia precursor .......... S9

6. $IC_{50}$ assay - Cell free Assay .......... S10

7. Molecular Docking .......... S11

8. Radiolabeling .......... S12

9. Biology: Methods ..... S15
9.1 Log P & Log D ..... S15
9.2 Plasma Stability ..... S15
9.3 Cell-free pulldown study from DNA ..... S16
9.4 Cell-free pulldown study from PARP ..... S16
9.5 PARP levels associated with chromatin ..... S18
9.6 Dosimetry calculation ..... S19
10. References ..... S19

#### 1. General

All chemicals were purchased from Sigma-Aldrich Co. and BLD Pharma and were used without further purification. Reactions were monitored using UPLC-MS via a Waters ACQUITY QDa Detector, and thin-layer chromatography (TLC) on 0.20 mm POLYGRAM SIL G/$UV_{254}$ (silica gel 60) TLC plates and were developed with an appropriate running buffer/solvent mixture. Spots were visualized with UV light (254 or 366 nm). $^{1}$H and $^{13}$C NMR spectra were obtained at 300 K using an Avance III AV 600 ($^{1}$H: 600.13 MHz and $^{13}$C: 150.61 MHz) spectrometer (Bruker Biospin, Ettlingen, Germany). All chemical shifts (δ) are reported in parts per million, and all *J* values are reported in Hz. The following abbreviations are used to describe multiplicities: s (singlet), d (doublet), t (triplet), q (quartet), m (multiplet), and brs (broad singlet). All compounds were dissolved in $CDCl_3$ or DMSO unless otherwise stated. All chemical shifts were referenced to residual chloroform ($\delta_H$ = 7.24 and $\delta_C$ = 77.00) or DMSO ($\delta_H$ = 2.50 and $\delta_C$ = 39.52). Analytical HPLC-MS data were collected using a Waters ACQUITY QDa Detector. Chromatographic purification was performed using an AKTA Purifier equipped with a UV-900 system, P-900 pump, frac-920 fraction collector, and a XTerra MS C18 OBD Prep Column, 125Å, 5 µm, 19 mm X 100 mm column. Solvent A: $H_2O$ + trifluoroacetic acid (0.1%); solvent B: acetonitrile; gradient: 0–8 CV (30% B), 8-10 CV (100% B), 10-14 CV (100% B), CV= column volume. Compounds **8**, **9** and **10** have been initially characterized only by MS (ESI+) and $^{1}$H NMR to later perform the PARP $IC_{50}$ assay purchased from BPS Bioscience, catalogue #80551, San Diego, CA, USA.

#### 2. Synthesis:

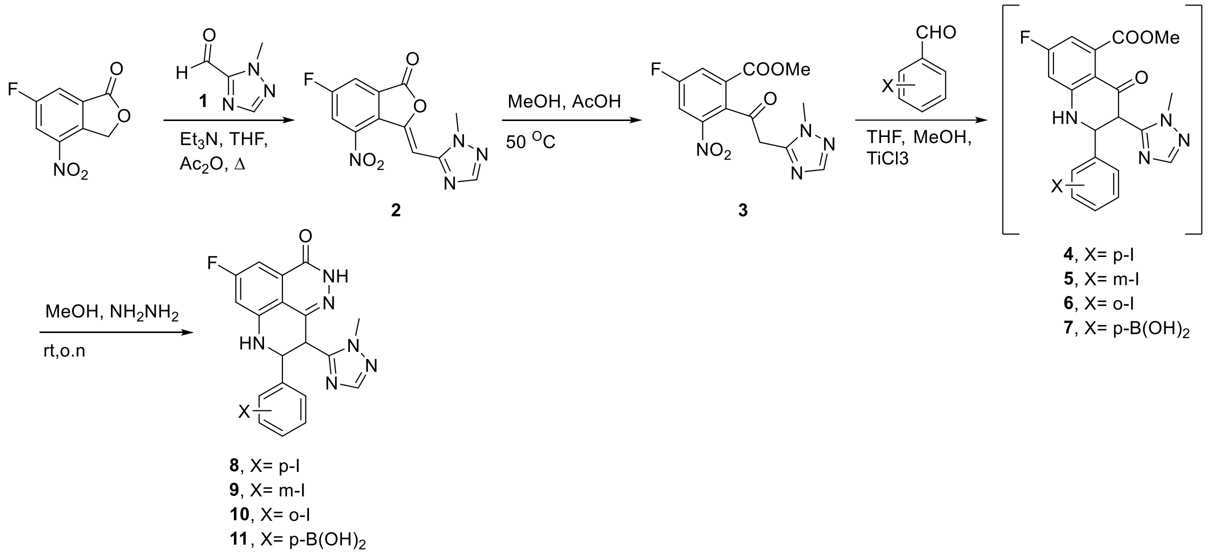


**Suppl. Scheme 1**: General synthesis of iodinated talazoparib derivatives (**8**, **9**, **10**) and boronic acid precursor (**11**). This synthetic pathway is adapted from G. Bowden *et al.*[1]

#### 3. Synthesis of compounds 8, 9, 10 and 11:

##### 3.1 5-fluoro-8-(4-iodophenyl)-9-(1-methyl-1H-1,2,4-triazol-5-yl)-2,7,8,9-tetrahydro-3H-pyrido[4,3,2-de]phthalazin-3-one (8)

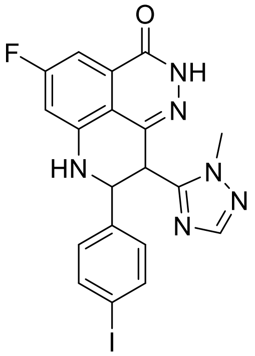

Chemical Formula: $C_{19}H_{14}FIN_6O$
Molecular Weight: 488.26
m/z: 489.03

Compound **3** (0.10 g, 0.031 mmol) and 4-iodobenzaldehyde (0.144 g, 0.062 mmol) were suspended in THF (0.6 mL) and MeOH (0.15 mL). To the resulting mixture was added titanium(III) chloride solution [20% wt solution in HCl (2 M), 1.6 mL, 6 equiv.] in dropwise fashion slowly at room temperature. The reaction temperature was maintained between 30 and 50°C for 2 h, after which it was quenched by the slow addition of water (3.2 mL). The reaction mixture was then poured into a separating funnel and extracted with ethyl acetate (4 × 20 mL). The organic fractions were pooled and washed with $NaHCO_3$ (3 × 10 mL) and $NaHSO_3$ (3 × 10 mL), dried with sodium sulphate ($Na_2SO_4$), and concentrated under reduced pressure to afford a think yellow syrup, which was carefully washed with aliquots of diethyl ether (3 × 10 mL). The resulting yellow syrup was then dried under high vacuum to afford the crude intermediate **4** as a yellow amorphous solid (0.152 g, 96%) that was used in the next step without any further purification. The resulting intermediate was dissolved in methanol (0.40 mL) at room temperature, and to the resulting solution was added hydrazine monohydrate (0.1 mL). The reaction mixture was then left to stir overnight at room temperature. After solvent evaporation, the crude was purified using an AKTA Purifier with the general method described above to yield the product **8** as a yellow powder (0.049g, 0.55mmol, 34% yield).

$^1$HNMR (500 MHz, $CD_3OD$): 7.89 (d, J = 1.6 Hz, 1H), 7.64 (d, J = 44.8 Hz, 2H), 7.38 (s, 2H), 7.30 – 7.13 (m, 1H), 6.91 (ddd, J = 10.9, 3.4, 2.0 Hz, 1H), 3.64 (s, 3H), 3.39 – 3.33 (m, 1H).

##### 3.2 5-fluoro-8-(3-iodophenyl)-9-(1-methyl-1H-1,2,4-triazol-5-yl)-2,7,8,9-tetrahydro-3H-pyrido[4,3,2-de]phthalazin-3-one (9)

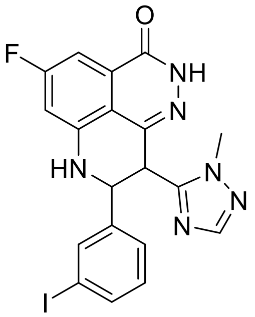


Chemical Formula: $C_{19}H_{14}FIN_6O$
Molecular Weight: 488.26
m/z:489.03 (20.7%),

Compound **3** (0.10 g, 0.031 mmol) and 3-iodobenzaldehyde (0.144 g, 0.062 mmol) were suspended in THF (0.6 mL) and MeOH (0.15 mL). To the resulting mixture was added titanium(III) chloride solution [20% wt solution in HCl (2 M), 1.6 mL, 6 equiv.] in dropwise fashion slowly at room temperature. The reaction temperature was maintained between 30 and 50°C for 2 h, after which it was quenched by the slow addition of water (3.2 mL). The reaction mixture was then poured into a separating funnel and extracted with ethyl acetate (4 × 20 mL). The organic fractions were pooled and washed with $NaHCO_3$ (3 × 10 mL) and $NaHSO_3$ (3 × 10 mL), dried with sodium sulphate ($Na_2SO_4$), and concentrated under reduced pressure to afford a think yellow syrup, which was carefully washed with aliquots of diethyl ether (3 × 10 mL). The resulting yellow syrup was then dried under high vacuum to afford the crude intermediate **5** as a yellow amorphous solid (0.150 g, 95%) that was used in the next step without any further purification. The resulting intermediate was dissolved in methanol (0.40 mL) at room temperature, and to the resulting solution was added hydrazine monohydrate (0.1 mL). The reaction mixture was then left to stir overnight at room temperature. After solvent evaporation, the crude was purified using an AKTA Purifier with the general method described above to yield the product **9** as a yellow powder (0.043 g, 0.48 mmol, 30% yield).

$^1$HNMR (600 MHz, DMSO): d 12.35 (s, 1H), 7.87 (s, 1H), 7.74 (s, 1H), 7.66 (dd, $J$ = 7.8, 1.7 Hz, 1H), 7.42 (d, $J$ = 7.6 Hz, 1H), 7.12 (d, $J$ = 7.8 Hz, 1H), 7.07 (dd, $J$ = 9.0, 2.5 Hz, 1H), 6.91 (dd, $J$ = 11.1, 2.5 Hz, 1H), 5.07-4.93 (m, 2H), 3.68 (s, 3H). MS (ESI): m/z calcd for $C_{19}H_{14}FIN_6O$ [M+H]+, 489.03; found, 489.033.

##### 3.3 5-fluoro-8-(2-iodophenyl)-9-(1-methyl-1H-1,2,4-triazol-5-yl)-2,7,8,9-tetrahydro-3H-pyrido[4,3,2-de]phthalazin-3-one (10)

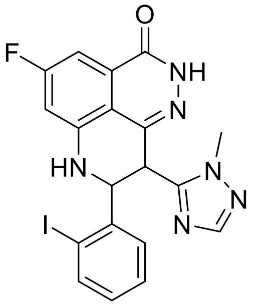

Chemical Formula: $C_{19}H_{14}FIN_6O$
Molecular Weight: 488.26
m/z:489.03 (20.7%),

Compound **3** (0.10 g, 0.031 mmol) and 2-iodobenzaldehyde (0.144 g, 0.062 mmol) were suspended in THF (0.6 mL) and MeOH (0.15 mL). To the resulting mixture was added titanium (III) chloride solution [20% wt. solution in HCl (2 M), 1.6 mL, 6 equiv.] in dropwise fashion slowly at room temperature. The reaction temperature was maintained between 30 and 50°C for 2 h, after which it was quenched by the slow addition of water (3.2 mL). The reaction mixture was then poured into a separating funnel and extracted with ethyl acetate (4 × 20 mL). The organic fractions were pooled and washed with $NaHCO_3$ (3 × 10 mL) and $NaHSO_3$ (3 × 10 mL), dried with sodium sulphate ($Na_2SO_4$), and concentrated under reduced pressure to afford a think yellow syrup, which was carefully washed with aliquots of diethyl ether (3 × 10 mL). The resulting yellow syrup was then dried under high vacuum to afford the crude intermediate **6** as a yellow amorphous solid (0.076 g, 48%) that was used in the next step without any further purification. The resulting intermediate was dissolved in methanol (0.20 mL) at room temperature, and to the resulting solution was added hydrazine monohydrate (0.05 mL). The reaction mixture was then left to stir overnight at room temperature. After solvent evaporation, the crude was purified using an AKTA Purifier with the general method described above to yield the product **10** as a yellow powder (0.021 g, 0.55 mmol, 30% yield).

$^1$HNMR (600 MHz, DMSO): d 12.38 (s, 1H), 7.84 (dd, $J$ = 7.9, 1.2 Hz, 1H), 7.76 (s, 1H), 7.73 (s, 1H) 7.69 (dd, $J$ = 8.0, 1.6 Hz, 1H), 7.42 (td, $J$ = 7.5, 1.3 Hz, 1H), 6.92 (dd, $J$ = 11.1, 2.5 Hz, 1H).5.28-5.20 (m, 2H), 3.77 (s, 3H). MS (ESI): m/z calcd for $C_{19}H_{14}FIN_6O$ [M+H]+, 489.03; found, 489.033.

##### 3.4 (4-(5-fluoro-9-(1-methyl-1H-1,2,4-triazol-5-yl)-3-oxo-2,7,8,9-tetrahydro-3H-pyrido[4,3,2-de]phthalazin-8-yl)phenyl)boronic acid (11)

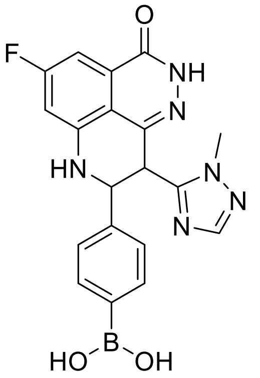

Chemical Formula: $C_{19}H_{16}BFN_6O_3$
Molecular Weight: 406.18
m/z: 405.14

Compound **3** (0.10 g, 0.031 mmol) and 4-boronic acid benzaldehyde (0.093 g, 0.062 mmol) were suspended in THF (0.6 mL) and MeOH (0.15 mL). To the resulting mixture was added titanium(III) chloride solution [20% wt. solution in HCl (2 M), 1.6 mL, 6 equiv.] in dropwise fashion slowly at room temperature. The reaction temperature was maintained between 30 and 50°C for 2 h, after which it was quenched by the slow addition of water (3.2 mL). The reaction mixture was then poured into a separating funnel and extracted with ethyl acetate (4 × 20 mL). The organic fractions were pooled and washed with $NaHCO_3$ (3 × 10 mL) and $NaHSO_3$ (3 × 10 mL), dried with sodium sulphate ($Na_2SO_4$), and concentrated under reduced pressure to afford a think yellow syrup, which was carefully washed with aliquots of diethyl ether (3 × 10 mL). The resulting yellow syrup was then dried under high vacuum to afford the crude intermediate **7** as a yellow amorphous solid (0.130 g, 96%) that was used in the next step without any further purification. The resulting intermediate was dissolved in methanol (0.40 mL) at room temperature, and to the resulting solution was added hydrazine monohydrate (0.1 mL). The reaction mixture was then left to stir overnight at room temperature. After solvent evaporation, the crude was purified using an AKTA Purifier with the general method described above to yield the product 11 as a yellow powder (0.049 g, 0.55 mmol, 34% yield).

$^1$HNMR (500 MHz, $CD_3OD$): 7.89 (d, *J* = 1.6 Hz, 1H), 7.64 (d, *J* = 44.8 Hz, 2H), 7.38 (s, 2H), 7.30 – 7.13 (m, 1H), 6.91 (ddd, *J* = 10.9, 3.4, 2.0 Hz, 1H), 3.54 (s, 3H), 3.39 – 3.33 (m, 1H). $^{13}$C NMR (500 MHz, $CD_3OD$) δ 154.09, 151.14, 143.74, 131.24, 128.10, 113.28, 105.12, 104.85, 100.92, 100.67, 62.25, 49.80, 49.66, 49.58, 49.44, 49.37, 49.16, 48.94, 48.73, 48.52, 45.65, 35.66. MS (ESI): m/z calcd for $C_{19}H_{16}BFN_6O_3$ [M] = 406.14; found, 407.23.

### 4. NMR Spectra

#### 4.1 $^1$HNMR Compound 8

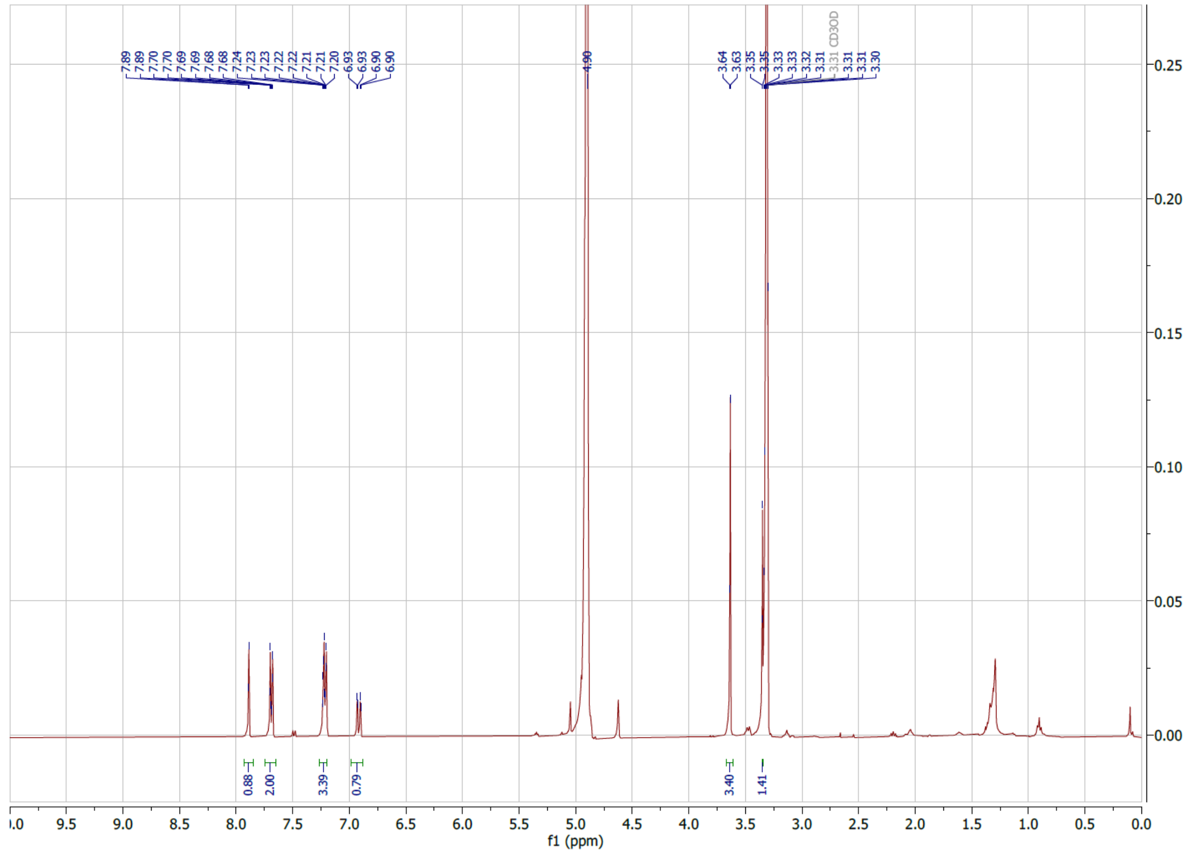

#### 4.2 [1]HNMR Compound 9

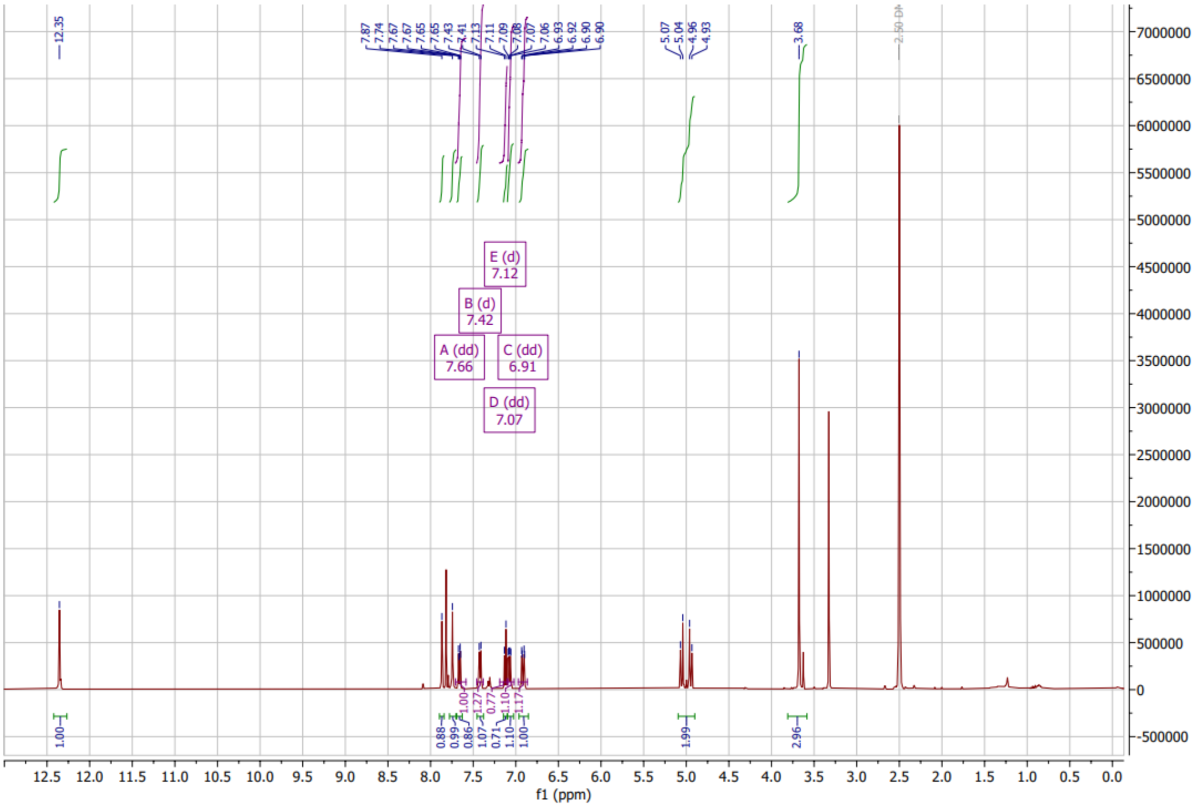

#### 4.3 1HNMR Compound 10

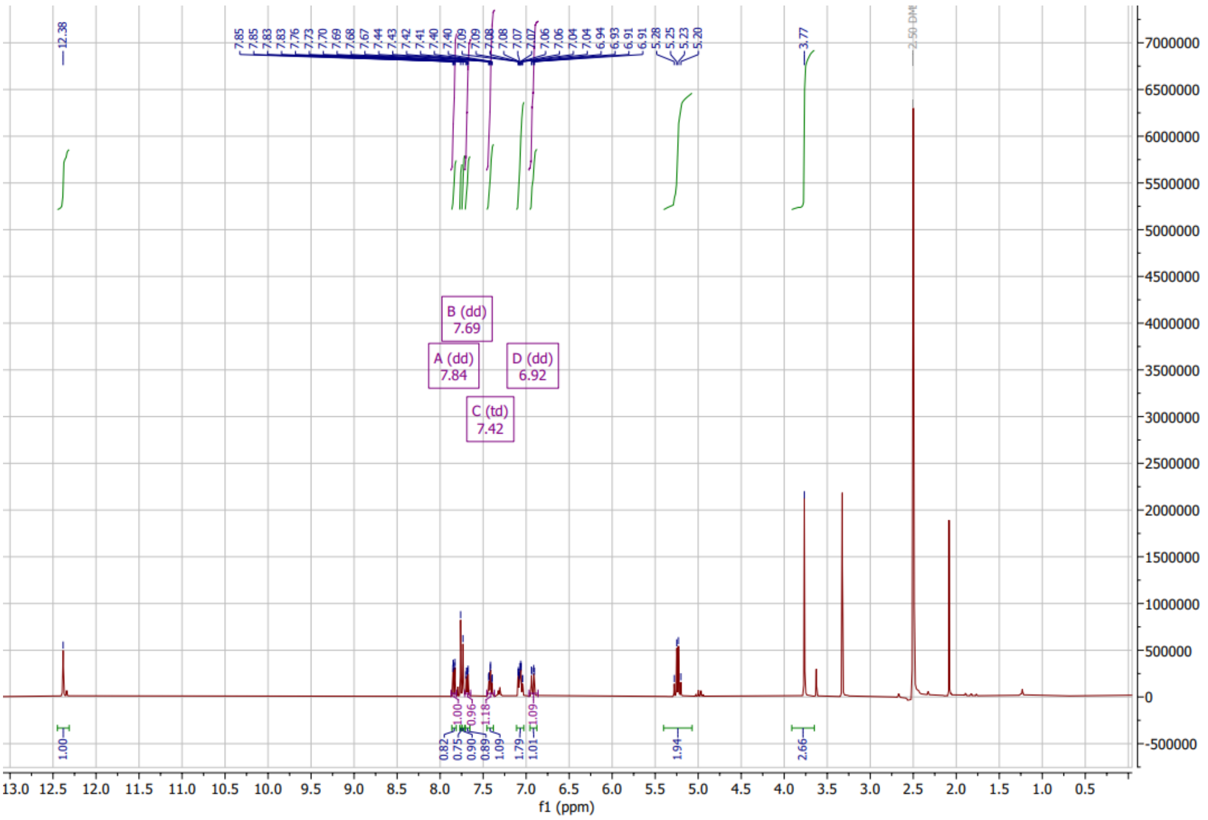


#### 4.4 $^{1}$HNMR Compound 11

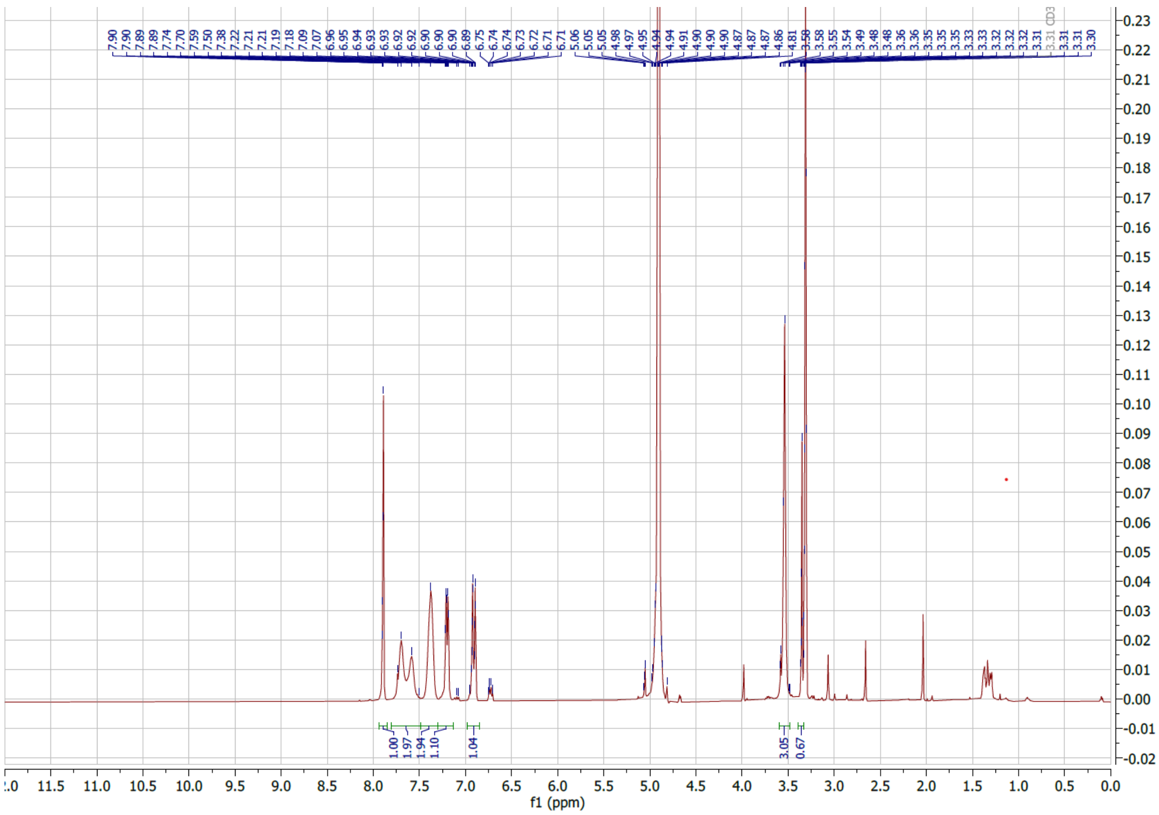

##### 4.5 $^{13}$CNMR Compound 11

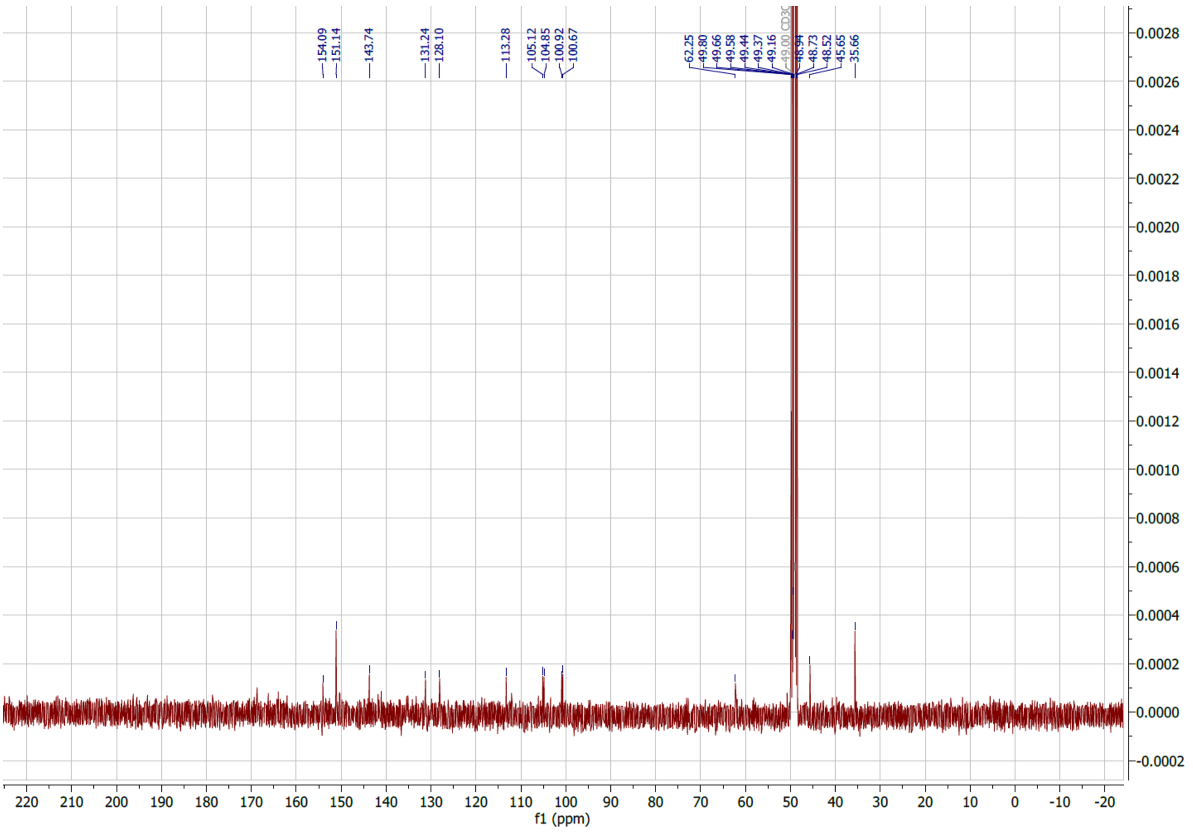


#### 5. Chiral HPLC separation of Italia precursor

Enantiomeric separation was performed on an ÄKTA FPLC system equipped with UV detection (λ = 280, 254, and 210 nm) using a CHIRALPAK® IB N-5 analytical column. The mobile phases consisted of (A) $H_2O$ containing 0.1% TFA and (B) MeCN. The flow rate was 0.5 mL/min with a system pressure limit of 25 MPa. The gradient was programmed from 10% to 30% B over 6 CV, followed by isocratic elution at 30% B to a total of 15 CV. Fractions were collected at 1.0 mL per tube. Under these conditions, enantiomer A eluted at 6.0-7.5 CV (30-38 min), and enantiomer B eluted at 7.6–9.0 CV (39-45 min) (**Figure S1**).

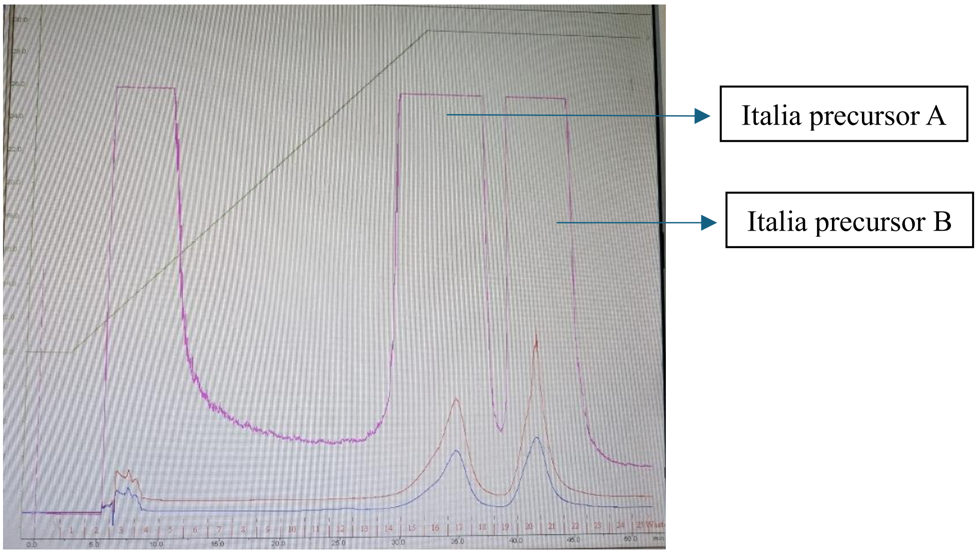


**Suppl. Figure S1**: Chiral HPLC separation of talazoparib on a CHIRALPAK® IB N-5 analytical column.

#### 6. $IC_{50}$ assay - Cell free Assay

A commercially available PARP-1 Chemiluminescent Assay Kit (BPS Bioscience, catalogue #80551, San Diego, CA, USA) was employed to measure catalytic inhibition of PARP activity by *p*-I-talazoparib (**8**), *m*-I-talazoparib (**9**), *o*-I-talazoparib (**10**) compared to talazoparib, following the manufacturer's instructions.

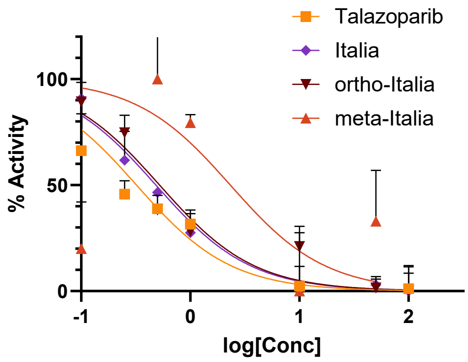


| Compound | $IC_{50}$ (nM) | R squared | 95% CI |
|---|---|---|---|
| Talazoparib | 0.32 | 0.8695 | 0.23 – 0.44 |
| Italia | 0.48 | 0.9334 | 0.37 – 0.63 |
| ortho-Italia | 0.52 | 0.8869 | 0.37 – 0.73 |
| meta-Italia | 2.35 | 0.2168 | 0.33 – 22.3 |

**Suppl. Figure S2**: **A)** Concentration-response curves for talazoparib, Italia, o-I-talazoparib, and m-I-talazoparib. **B) B)** Cell-free enzymatic inhibition of PARP1 by Italia (**8**), ortho-Italia (**10**), meta-Italia (**9**) or talazoparib. $IC_{50}$ = inhibitory concentration of 50%.

#### 7. Molecular Docking

The solved X-ray crystal structure (PDB code: 7KK3)[2] of the catalytic domain of PARP1 in complex with talazoparib, with a resolution of 2.06 Å was selected as a reference point for the molecular docking studies. Structures of the PARP1, PARP2 and PARP3 were obtained from AlphaFold[3] (AF-P09874-F1, AF-Q9UGN5-F1-v4 and AF-Q9Y6F1-F1-v4) and aligned with the reference PDB file. AutoDock Tools 1.5.7[4] was used to perform docking studies, providing predicted binding poses and binding energies for the ligands interacting with the receptor. All ligands were docked to the rigid receptor. Binding sites were specified in the same way as the crystal structure. Grid box was chosen 70×70×70 points with a spacing of 0.375 Å. For all dockings, docking parameters were used in their default values. All interactions between ligand and receptor were visualized using PYMOL software[5] for 3D diagrams and Discovery Studio Visualizer[6] for 2D diagrams. Italia was compared to the reference compound (F-)talazoparib based on the binding energies (kcal/mol) of their highest-scoring docking poses, and the resulting conformations were superimposed for comparison. **(Figure S3)**.

**PARP1**

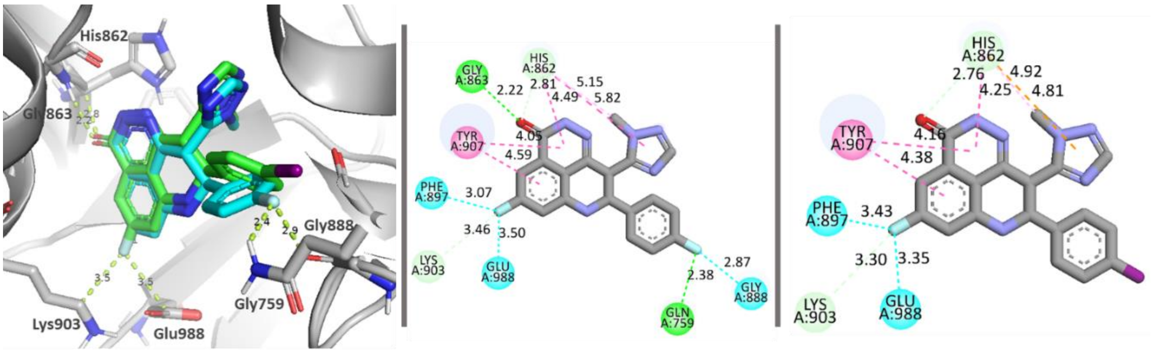


**PARP2**

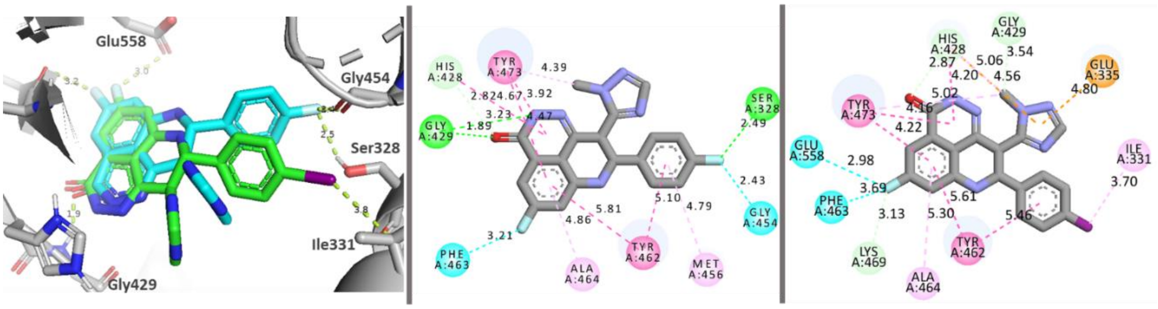


**PARP3**

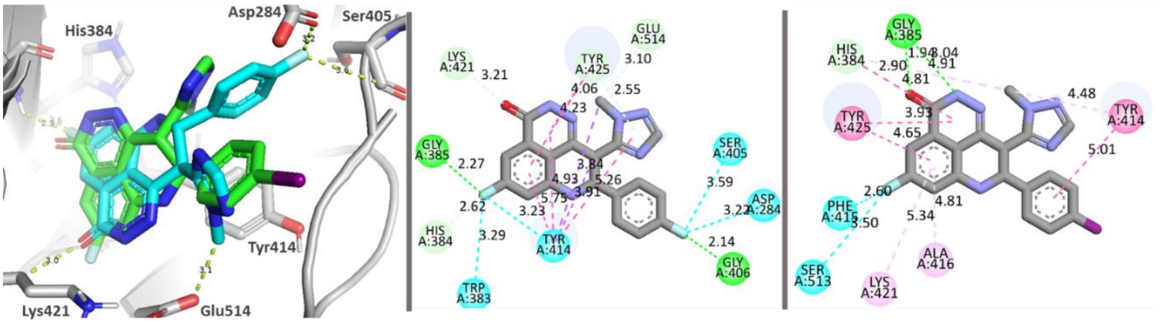


**Suppl. Figure S3a: A)** Comparison of docking of Italia (green) to talazoparib (blue) to PARP1, PARP2, and PARP3 and their interactions. **B)** Comparison of docking of Italia (green) to talazoparib (blue) to PARP1, PARP2, and PARP3 and their interactions. **C)** Comparison of docking of Italia (green) to talazoparib (blue) to PARP1, PARP2, and PARP3 and their interactions.

Comparison of AlphaFold docking to the crystal structure docking, with talazoparib:

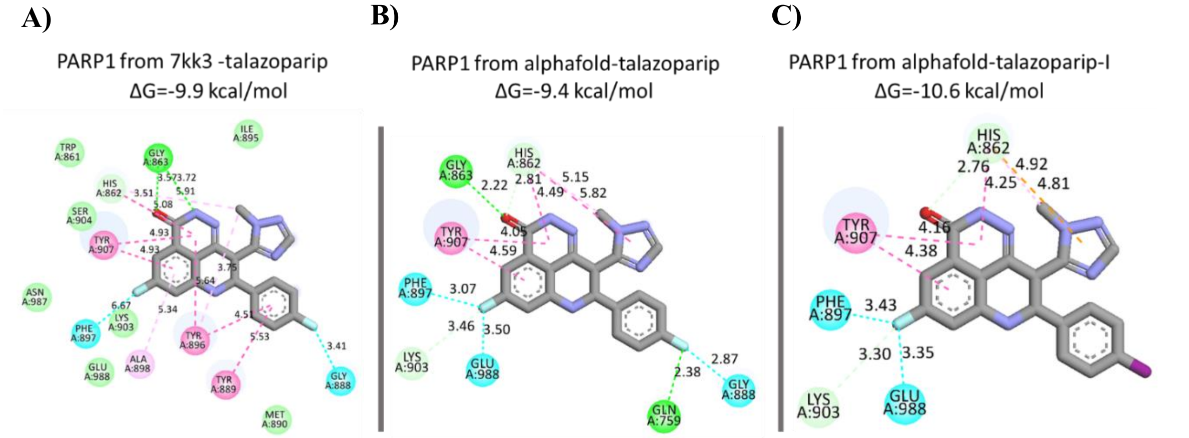


**Suppl. Figure S3b: A)** Docking performed using the crystal structure of PARP1 (PDB ID: 7KK3). **B)** Docking of talazoparib into the AlphaFold-predicted structure of PARP1. **C)** Docking of Italia to PARP1 and analysis of their interactions.

#### 8. Radiolabelling

Na[$^{123}$I]I was obtained from GE Healthcare as a non-carrier-added formulation in 0.05 M NaOH (typically delivered as a 18.7±1.4 GBq/mL solution). HPLC purification was conducted using an Elite LaChrom VWR Hitachi L-2130 pump and a Waters Symmetry C18 column. Detection was achieved with a UV detector (Elite LaChrom VWR Hitachi L-2400) set at 254 nm and a Bicron Frisk-Tech radiation detector. The mobile phase consisted of 30% acetonitrile (ACN) in water, delivered isocratically at a flow rate of 2 mL/min.

Radiolabelling was performed by treatment of Na[$^{123}$I] with the corresponding boronic acid precursor (0.1 mg) in the presence of copper catalysts Cu[(Ph-py)$_4$(ClO$_4$)]$_2$ (0.1 mg). The reaction was carried out in MeOH/$H_2O$ (4:1) for 25 min. After completion, the mixture was diluted with MeCN/$H_2O$ (1:1) to 1 mL and filtered through a 0.2 µm hydrophobic PTFE filter. Radioactivity measurements were performed before and after filtration to ensure quantitative transfer.

The crude product was purified by semi-preparative reversed-phase HPLC (Luna Omega Polar-C18, 100 Å, 250 × 10 mm) using 30% acetonitrile in water at a flow rate of 2 mL/min. The radioactive product fraction was collected, diluted with water, and trapped on an HLB Plus solid-phase extraction cartridge. After washing with water, the product was eluted with ethanol. Radioactivity was monitored throughout purification to confirm maximal recovery.

The solvent was removed under gentle heating (80°C), and the residue was reconstituted in 10% DMSO/$H_2O$ (0.5-1 mL) to afford the final [$^{123}$I]Italia formulation.

**A)**

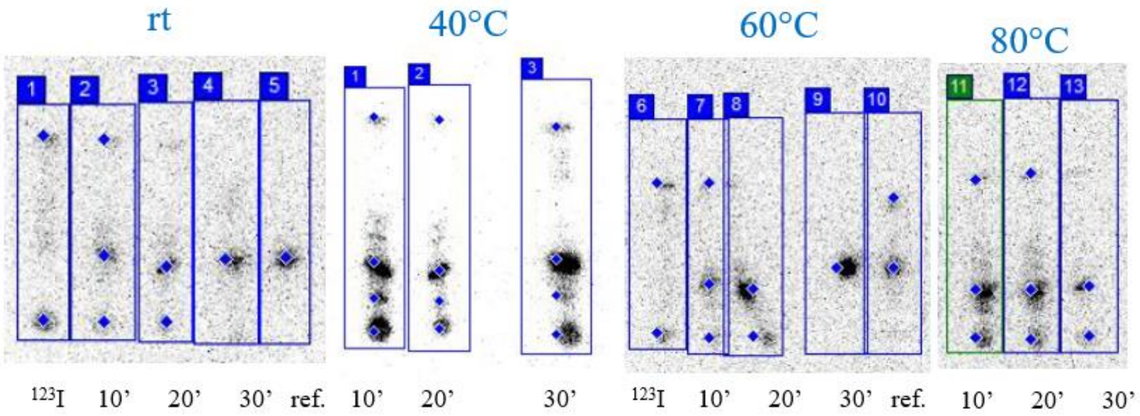


| Temperature °C | RCY(10 min) (%) | RCY(20 min) (%) | RCY(30 min) (%) |
|---|---|---|---|
| rt | 53 | 72 | 100 |
| 40 | 49 | 56 | 58 |
| 60 | 54 | 72 | 100 |
| 80 | 68 | 64 | 61 |

**B)**

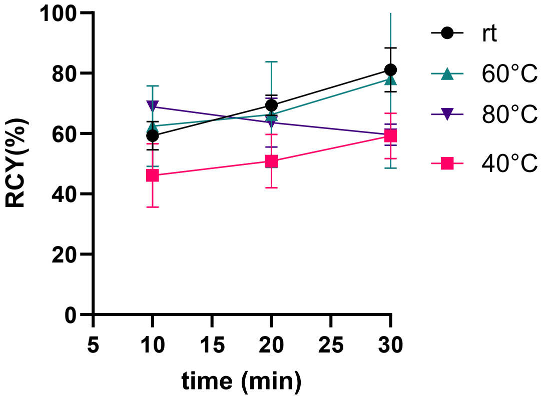


**Suppl. Figure S4**: Radiosynthesis [$^{123}$I]Italia: **A)** Optimization of labelling conditions at different temperatures – radio TLC analysis (eluted in 5% MeOH/DCM). **B)** Graphical representation of optimization of labelling conditions at different temperatures.

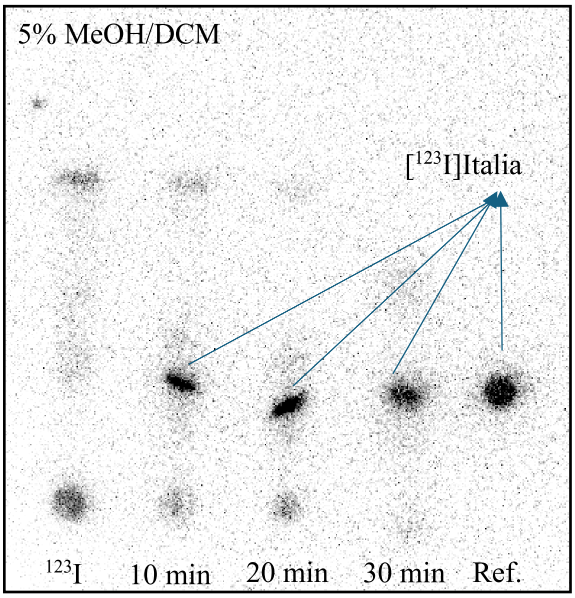


**Suppl. Figure S5**: Radio TLC of crude reaction mixture (RM) of radiosynthesis of [$^{123}$I]Italia.

The molar activity of [$^{123}$I]Italia was assessed by radiochromatography, using an ACQUITY UPLC® BEH SHIELD RP18 1.7 µm (186004668) column eluted with 30% ACN/water over 12 min (0.6 mL/min) monitoring with UV (254 nm) and radioactive traces, with concentration 1 mg/mL as LOQ.

**A)**

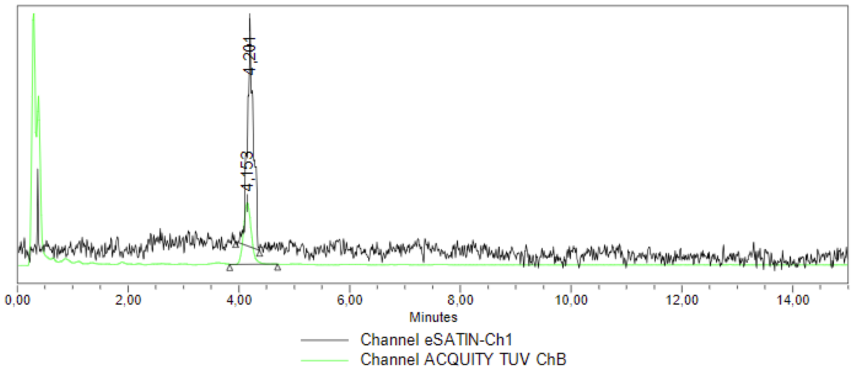


**B)**

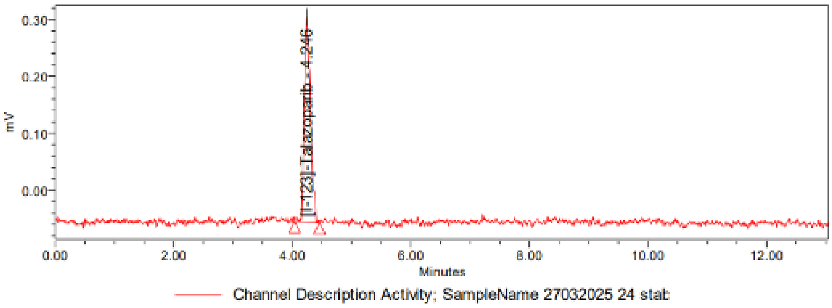


**Suppl. Figure S6**: **A)** Radiochromatogram obtained during the chromatography analysis of [$^{123}$I]Italia (black) at 0 h overlayed with the UV chromatogram (green) of the same sample. **B)** Radiochromatogram obtained during the chromatography analysis of [$^{123}$I]Italia at 24 h.

#### 9. Biology: Methods

##### 9.1 Log P & Log D

[$^{123}$I]Italia (10 µL, 10 kBq) was added to two separate biphasic systems prepared in Eppendorf tubes: (a) PBS/octanol (1:1, v/v, Log D) and (b) water/octanol (1:1, v/v, Log P). Each mixture was vortexed for 1 min and then incubated at room temperature for 10 min to allow phase separation. After incubation, the octanol layer was carefully collected and transferred into a counting tube. The aqueous layers (PBS or water) were collected separately into corresponding counting tubes. All samples were then measured using a gamma counter.

##### 9.2 Plasma Stability

[$^{123}$I]Italia (100 kBq) was added to 1 mL of sterile-filtered human male AB plasma (Sigma-Aldrich) in Eppendorf tubes. Samples were prepared in at least triplicate for each time point. The mixtures were vortexed for 1 min and incubated at 37°C for 15 min, 1 h, 3 h, 6 h, and 24 h. At each designated time point, a 2 µL aliquot was withdrawn and applied to a silica TLC plate. The plates were developed in 5% MeOH/DCM and analysed using a Typhoon phosphor imager to assess radiochemical stability.

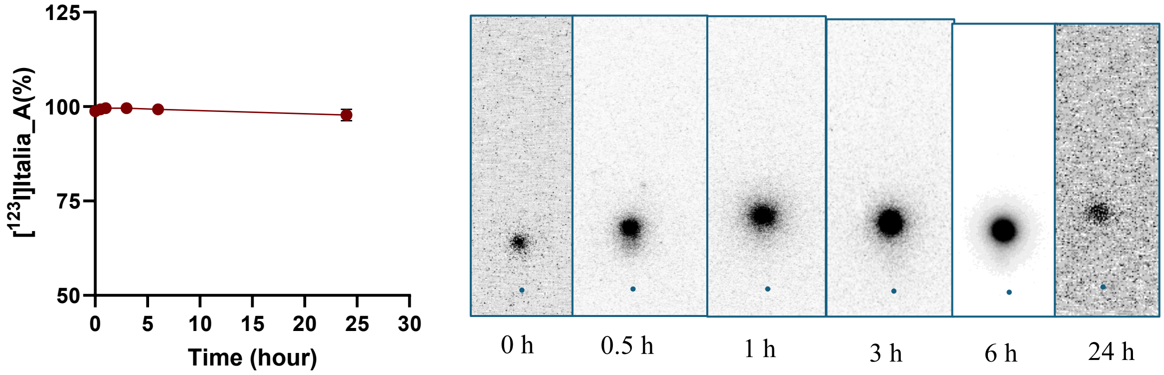


**Suppl. Figure S7**: Human plasma stability of [$^{123}$I]Italia assessed by radio-TLC.

#### 9.3 Cell-free pulldown study from DNA

Reactions were carried out in pull-down buffer containing: 50 mM Tris-HCl (pH 8.0), 10 mM $MgCl_2$, 1 mM DTT, and 10% (v/v) glycerol. Mixtures were prepared in Eppendorf tubes and contained the following components according to experimental conditions: 50 nm recombinant PARP (Medchem, HY-P74652), Biotinylated synthetic damaged DNA (containing one single strand break), where indicated, olaparib (10 µM final) and/or [$^{123}$I]Italia (100 kBq) in a total volume of 150 µL. Samples were incubated at 30°C for 1 h, followed by cooling on ice to terminate the reaction. A small aliquot of each sample was retained for total protein quantification or activity quantification prior to pull-down. For each sample, 25 µL of streptavidin-agarose bead slurry was used. Beads were washed at least three times with pull-down buffer to remove ethanol, then blocked with 5% BSA in pull-down buffer for 60 minutes at 4°C. After blocking, beads were washed again three times and resuspended in pull-down buffer. Each 200 µL reaction was brought to 150 µL total volume with pull-down buffer containing 1% BSA, and resuspended beads were added. Samples were incubated for 1 hour on a rotating wheel at 4°C. Beads were then washed three times with 0.5-1 mL of pull-down buffer, with centrifugation at 2500 × g for 2 min to pellet the resin. Supernatants were removed each time. Finally, resin-bound complexes were boiled in SDS sample buffer, centrifuged, and the supernatant was analysed by western blot or gamma counting to quantify PARP and [$^{123}$I]Italia binding.

#### 9.4 Cell-free pulldown study from PARP

Single strand Break constructs were incubated with His-Tagged Recombinant Human PARP (Medchem, HY-P74652) in the presence or absence of olaparib (10 mM), talazoparib (10 mM) or [$^{123}$I]Ital*i*a (100 kBq) for 1 hour at 30°C under constant agitation in buffer (50 mM Tris-HCl pH 8, 10 mM $MgCl_2$, 10 mM DTT and 10 % Glycerol). Streptavidin pulldown using Thermo Scientific™ Pierce™ Streptavidin Agarose (according to the manufacturer's instructions) was used to isolate btDNA and $^{123}$I content in fractions was measured using a gamma counter. To detect [$^{123}$I]Italia bound to PARP, the mixture was incubated with anti his-Tag (D3I1O) antibody (1:50, 12698, Cell Signaling) for one hour at 4°C under gentle agitation, followed by incubation with Protein G Agarose Resin 4 Rapid Run™ (ABT, 4RRPG-5) according to manufacturer's instructions. Pulldown fractions were collected for the quantification of $^{123}$I using a gamma-counter.

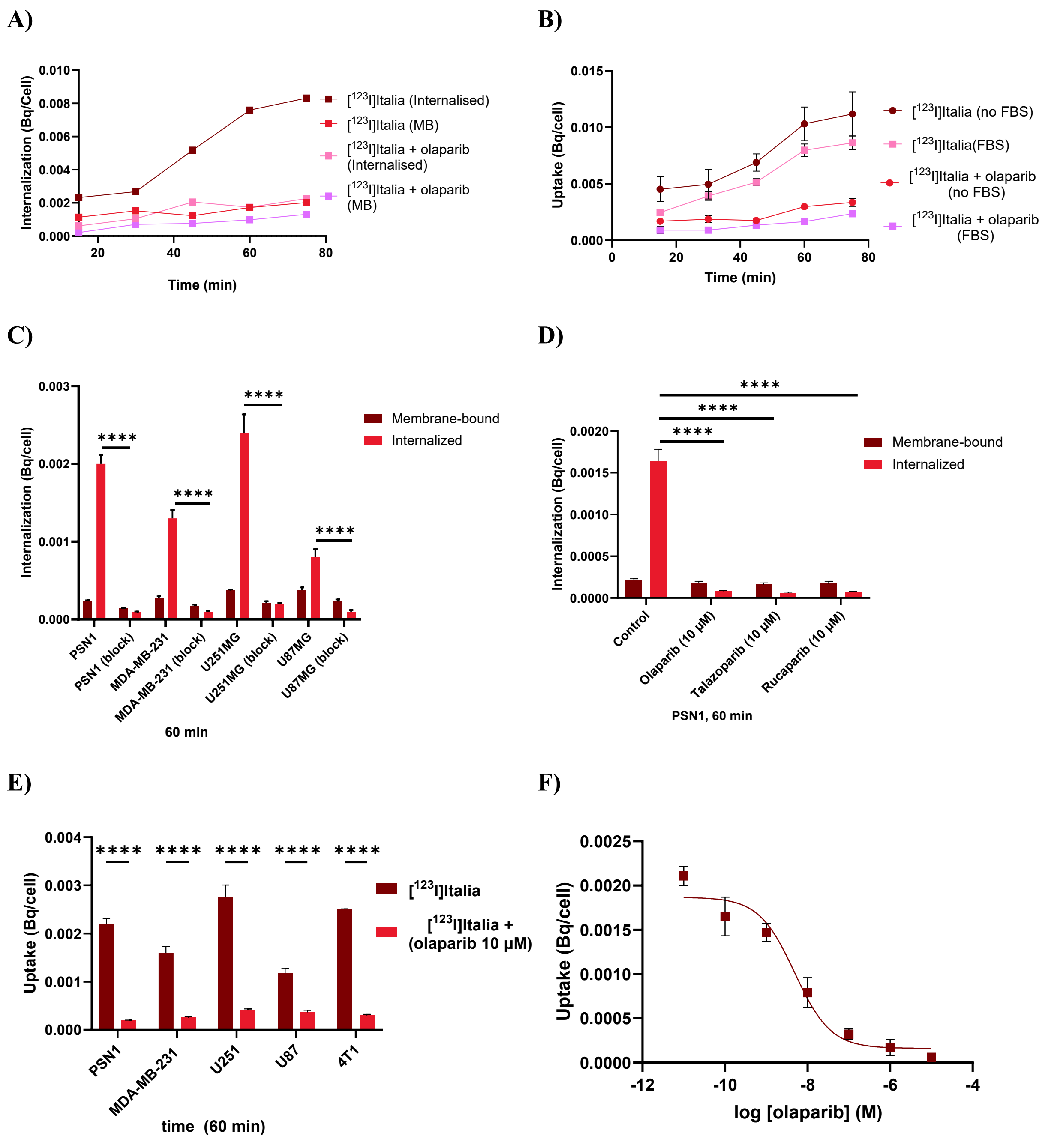


G)

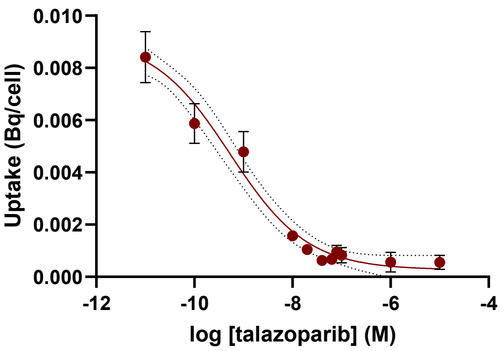


**Suppl. Figure S8: A)** Internalization of [$^{123}$I]Italia in PSN1 at different (shorter) time points. **B)** Uptake of [$^{123}$I]Italia in PSN1 with and without in FBS. **C)** Internalization and blocking of [$^{123}$I]Italia in PSN1, MSA-MB-231, U251, and U87 cells after 1 hour exposure. **D)** Blocking of [$^{123}$I]Italia uptake in PSN1 cells by panel of PARP inhibitors. **E)** Uptake and blocking of [$^{123}$I]Italia in PSN1, MDA-MB-231, U251, U87, and 4T1 cells after 1 hour exposure. **F&G)** IC50 in PSN1 cells (different concentration of olaparib (IC50 : 4.9 nM, 95% CI: 2.26 nM-9.82 nM, R squared: 0.9497 and talazoparib (IC50 = 0.53 nM, 95% CI: 0.2 nM-1.3 nM, R squared: 0.9599).

##### 9.5 PARP levels associated with chromatin

Aliquots of 5 x $10^6$ suspended cells were incubated with 200 kBq/mL of [$^{123}$I]Italia in a final volume of 500 µL after exposure to the specified drug treatments. The reaction was stopped on ice and cells were washed with ice-cold PBS. Cells were lysed in 9 M urea buffer (urea 9 M, tris-Base pH 7.5 1 M, 0.1% β-mercaptoethanol) supplemented with PhosSTOP (Merck, 4906845001) and cOmplete™, Mini, EDTA-free Protease Inhibitor Cocktail (Merck, 11836170001 ). To ensure the dissociation of proteins from the DNA, samples were sonicated (1 pulse, 2 seconds).

To determine PARP levels using western bolt, protein samples (20-40 mg) of chromatin and the nuclear soluble fraction, collected as described above, were used. Samples were loaded onto Bolt™ Bis-Tris Plus Mini Protein Gels (4-12%, 1.0 mm, Invitrogen™, Wedge Well™ format) and separated by SDS-PAGE at constant voltage. Proteins were transferred into PVDF membranes using the Mini Blot Module (Thermo Scientific). Membranes were blocked using 5% non-fat milk at room temperature for 1 h and incubated overnight at 4°C with an anti-PARP1 antibody (1:1000, Merck, HPA045168), followed by incubation with an HRP-conjugated rabbit IgG secondary antibody (1:5000, 50 min, RT, R&D, HAF008). Chemiluminescent signals were visualized using Thermo Scientific™ Pierce™ ECL Western Blotting Substrate and quantified using Fiji/ImageJ.

##### 9.6 Dosimetry calculation

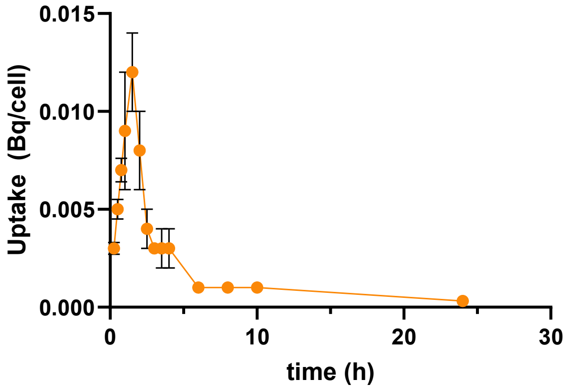


| | | | |
|---|---|---|---|
| Membrane | 7.304E-04 | 24.38 | 24.38 |
| Cytoplasm | 3.262E-04 | 10.89 | 10.89 |
| Nuclear | 1.573E-03 | 52.52 | 64.73 |
| Chr | 3.659E-04 | 12.21 | |
| Total | 2.996E-03 | | 100.00 |
| | For dosimetry calculations (uptake in Bq/cell in cell fractionation) | | |

**Area under the curve**: 143.064 Bq.s/cell

**S-value (Gy/Bq·s)** (calculated using MIRDcell): $3.1942 \times 10^{-3}$ Gy/Bq.s

(In PSN-1 cells, the distribution of [$^{123}$I]Italia_A determined by cell fractionation was 24% in the membrane-bound fraction, 11% in the cytoplasmic fraction, and 65% in the nuclear soluble fraction.)

Absorbed Dose to Nucleus, $\mathbf{D_N = A \times S_{total}}$

$\mathbf{D_N}$ = 0.46 Gy/cell (over 24 h)

5. Visualizing science with PyMOL 3 - Schrödinger. https://www.schrodinger.com/life-science/learn/education/courses/pymol/.

6. Dassault Systèmes’ Software Portfolio_ Unified by the 3DEXPERIENCE Platform - Dassault Systèmes.
